# Spatially resolved mapping of monoacylglycerol lipase activity in the brain

**DOI:** 10.1101/2025.07.08.663730

**Authors:** Daan van der Vliet, Alex X.Y. Klinkenberg, Rik Platte, Kieran Higgins, Susanne Prokop, Mirjam C.W. Huizenga, Lars Kraaijevanger, Noëlle van Egmond, Verena M. Straub, Maarten H. P. Kole, Pal Pacher, István Katona, Inge Huitinga, Mario van der Stelt

**Affiliations:** Department of Molecular Physiology, Leiden University, 2333 CC Leiden, The Netherlands; Department of Neuroimmunology, Netherlands Institute for Neuroscience, Institute of the Royal Netherlands Academy of Arts and Sciences, 1105 BA Amsterdam, The Netherlands; Department of Axonal Signaling, Netherlands Institute for Neuroscience, Institute of the Royal Netherlands Academy of Arts and Sciences, 1105 BA Amsterdam, The Netherlands; Cell biology, Neurobiology and Biophysics, Department of Biology, Faculty of Science, Utrecht University, Utrecht, The Netherlands; Momentum Laboratory of Molecular Neurobiology, Institute of Experimental Medicine, Budapest, Hungary; Department of Psychological and Brain Sciences, Indiana University, Bloomington, IN, USA; Laboratory of Cardiovascular Physiology and Tissue Injury, National Institute of Health/NIAAA, Rockville, MD, USA; Swammerdam Institute for Life Sciences, University of Amsterdam, Amsterdam, The Netherlands

**Keywords:** monoacylglycerol lipase, endocannabinoid, activity-based probe, hippocampus, histology, fluorescence, microscopy

## Abstract

Visualizing signaling systems in the brain with high spatial resolution is critical to understand brain function and to develop therapeutics. Especially enzymes are often regulated on the post-translational level, resulting in a disconnect between protein levels and activity. Conventional antibody-based methods have limitations, including potential cross reactivity and the inability of antibodies to discriminate between active and inactive enzyme states. Monoacylglycerol lipase (MAGL), an enzyme degrading the neuroprotective endocannabinoid 2-arachidonoylglycerol, is the target of inhibitors currently in clinical trials for the treatment of several neurological disorders. To support translational and (pre)clinical studies and fully realize the therapeutic opportunities of MAGL inhibitors, it is essential to map the spatial distribution of MAGL activity throughout the brain in both health and disease. Here, we introduce selective fluorescent activity-based probes for MAGL enabling direct visualization of its enzymatic activity in lysates, cultured cells and tissue sections. We show that oxidative stress, which inactivates MAGL through the oxidation of regulatory cysteines, reduces probe labeling, thereby validating the probes activity-dependence. Extending this approach, we developed an *activity-based histology* protocol to visualize MAGL activity in fresh-frozen mouse and human brain tissues. This approach revealed robust MAGL activity in astrocytes and presynaptic terminals within the mouse hippocampus, and further allows detection of MAGL activity in the human cerebral cortex. Collectively, these findings establish selective activity-based probes as powerful tools mapping MAGL activity with high spatial resolution across mammalian brain tissue.

## Introduction

The ability to visualize components of signaling systems in the brain with high spatial resolution is critical for understanding neurophysiological and pathophysiological processes.^1^ Traditional approaches in neuroscience, such as antibody-based immunohistochemistry or immunofluorescence, and gold particle-enhanced electron microscopy, remain widely used. However, these methods cannot distinguish between active and inactive protein states.^2,3^ This distinction is critical, as many signaling pathways are governed by post-translational regulation, making protein abundance a poor proxy for functional activity. This is especially true for enzymes, whose activities are often tightly regulated by substrate availability, protein-protein interactions, post-translational modifications and cofactor binding.

Pharmacological ligands that covalently interact with the active site residues of enzymes, also called activity-based probes (ABPs), can be used to map functional enzymes in the brain.^4–7^ These ABPs carry mechanism-based electrophiles that react with catalytic nucleophilic residues when enzymes are in an active state. Doing so, the enzyme is labeled by a reporter group, such as a fluorophore, that allows detection of the protein-probe adduct. Since the invention of ABPs for serine hydrolases 25 years ago^3^, these probes have been instrumental to guide inhibitor development and mapping of functional enzymes throughout the brain.^4,6,8–11^ However, the application of ABPs to map anatomical enzyme activity at high spatial resolution—referred to here as *activity-based histology*—remains limited. Most existing ABPs are broad-spectrum probes, targeting many enzymes simultaneously, thus requiring homogenization of tissues for analysis by gel electrophoresis or chemical proteomics.^8,12^ Furthermore, while imaging with activity-based probes has previously been reported,^13–16^ there is a need for additional robust protocols that preserve both enzyme activity and tissue architecture during labeling.

A major challenge in developing activity-based histology protocols for the brain lies in delivering probes to their enzymatic targets while preserving both activity and tissue integrity. This can be achieved through *in vivo* administration of fluorescent probes directly into the brain.^16^ More recently, Pang et al. described an approach using in vivo delivery of probes bearing bioorthogonal “click” handles, which are visualized post hoc via ligation to fluorescent tags.^17^ However, both these methods are labor intensive, require living animal experiments, and are not applicable to human tissue. Most human brain samples are obtained postmortem and stored either as formalin-fixed paraffin-embedded (FFPE) tissue or fresh-frozen material at −80 °C. Since fixation and embedding can compromise enzymatic activity, developing protocols for labeling active enzymes in cryosections from fresh-frozen tissue is of high translational value.

Monoacylglycerol lipase (MAGL) is an enzyme of the serine hydrolase family, whose inhibition has therapeutic potential in neurological disorders. MAGL metabolizes mono-acyl glycerols (MAGs), including 2-arachidonoylglycerol (2-AG), a central endocannabinoid.^18^ 2-AG regulates neurotransmission by activating cannabinoid receptor CB_1_, thereby providing a retrograde feedback signal to presynaptic terminals.^19,20^ Inhibition of neurotransmitter release finetunes neural circuits and avoids excessive neurotransmission as observed in epileptic seizures and excitotoxicity.^21,22^

2-AG also regulates glial cell biology, including microglial polarization and oligodendrocyte differentiation via CB_2_ and CB_1_ receptors, respectively.^23,24^ In addition, hydrolysis of 2-AG by MAGL is a rate limiting step for the supply of free arachidonic acid (AA) in the brain and liver which can be converted to pro-inflammatory prostaglandins, leukotrienes and thromboxanes.^25,26^ Thus, MAGL regulates both endocannabinoid and eicosanoid signalling.^27^ Blockade of MAGL ameliorates pathology and symptoms of mouse models of neuroinflammation, including models for Alzheimer’s Disease (AD)^28,29^, Multiple Sclerosis^30–34^ (MS) and Parkinson’s Disease (PD).^25^ Several MAGL inhibitors are currently in development as candidate therapeutics^35–37^, and recently ABX-1431 (Lu-AG06466 or Elcubragistat) has entered phase II clinical trials for several neurological disorders, such as spasticity in multiple sclerosis, Tourette Syndrome, post-traumatic stress disorder and neuromyelitis optica.^37^ Knowledge on how MAGL activity is distributed throughout the brain in health and disease would support the therapeutic development of MAGL inhibitors for CNS disorders.

We and others have recently developed selective fluorescent activity-based probes targeting MAGL in cells.^2,38–40^ Here, activity-based probes targeting MAGL were employed to image the localization of MAGL activity with high spatial resolution. The high selectivity of these probes allowed us to visualize MAGL activity in immortalized living cell lines and primary cortical cultures. Building on this, we established a protocol for activity-based histology to label MAGL activity in fresh-frozen mouse brain sections, allowing detailed neuroanatomical mapping in the hippocampus. Finally, we extended this approach to human brain tissue, demonstrating probe-based detection of MAGL activity in postmortem cortical sections. Together, these results establish activity-based probes as powerful tools for histological analysis of enzyme activity, with broad utility in translational and clinical neuroscience.

## Results

### Characterization of a set of fluorescent MAGL probes

MAGL is covalently targeted by a wide variety of activated carbamate inhibitors with diverse leaving groups (Figure 1a).^38,41^ Especially the hexafluoroisopropanol (HFIP) carbamate was shown to target MAGL with excellent selectivity.^38^ The HFIP carbamate is a bioisostere of the natural substrate of MAGL, the glycerol ester, and this moiety is incorporated in ABX-1431, a clinically safe MAGL inhibitor that is currently in phase II clinical trials.^35,37^ We synthesized an activity based probe bearing a piperazine coupled HFIP carbamate, functionalized with an azide ligation handle (probe **1**, Figure 1b)^2^. Subsequently three activity-based probes targeting MAGL were produced from **1**, equipped with three different fluorescent groups (Figure 1b).

**Figure 1.**
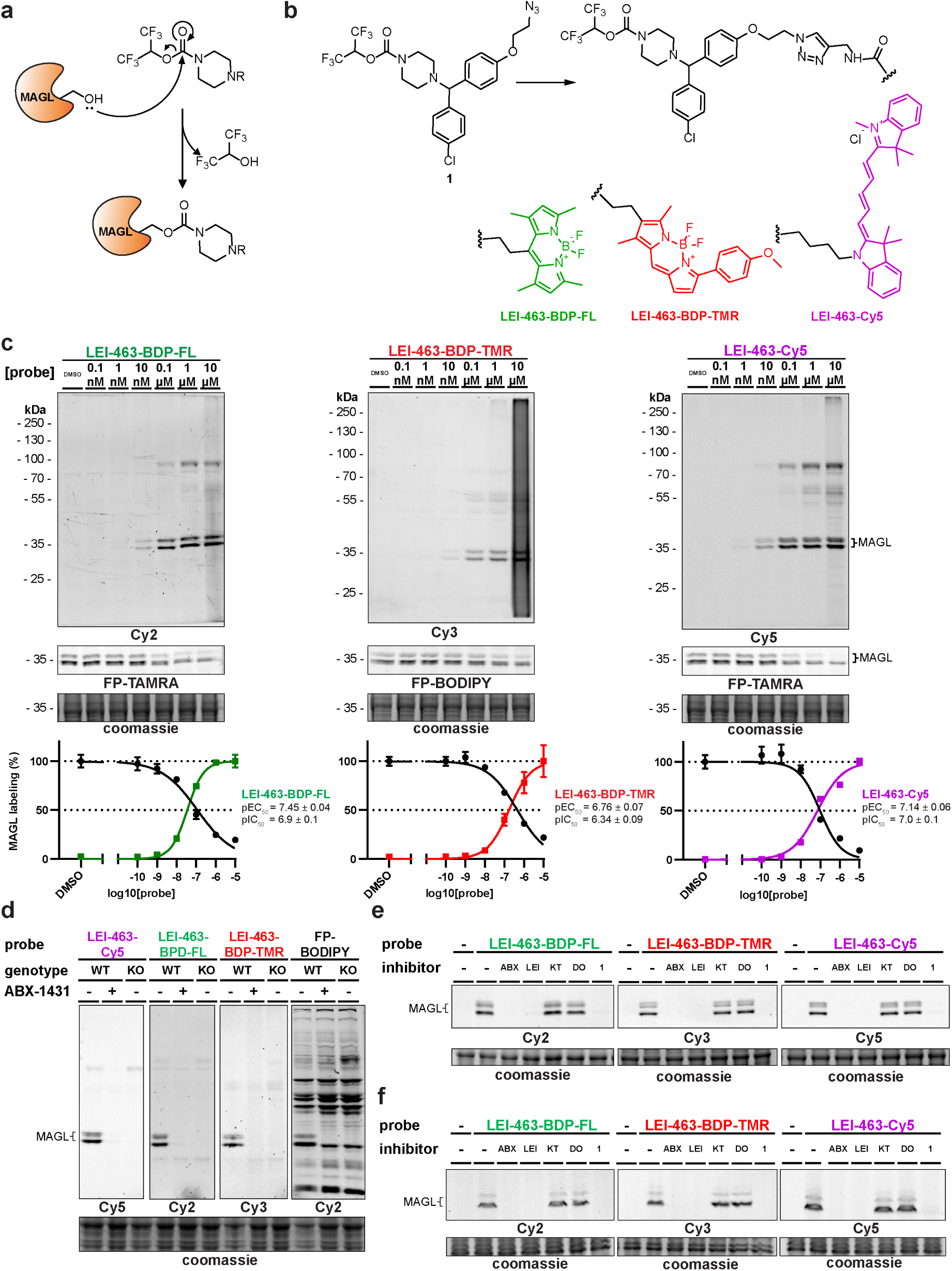
Selective and effective targeting of MAGL in mouse and human brain lysates. **a,** Mechanism of MAGL labeling by hexafluoroisopropanol-carbamates. **b,** Structures of the three probes synthesized in this study. The three probes bear the same MAGL targeting element, but are equipped with fluorophores with different excitation/emission wavelengths: **LEI-463-BDP-FL**, 493/503 nm; **LEI-463-BDP-TMR**, 542/574 nm, **LEI-463-Cy5**, 646/662 nm. **c,** Dose response labeling of MAGL in mouse brain proteome by the indicated **LEI-463** probes, and quantification of IC_50_ and EC_50_ values (N=3)**. d,** Absence of MAGL labeling after incubation with probe (100 nM, 30 min, RT) by either pre-incubation with ABX-1431 (10 µM) or in lysates from Mgll /- mice (indicated as KO). **e-f,** Mouse brain lysates (**e**) or human cortical lysates (**f**) were incubated with the indicated inhibitors prior to treatment with the **LEI-463** probes (100 nM). Abbreviations: ABX, ABX-1431; LEI, LEI-515; KT, KT182; DO, DO264.

To assess efficacy and selectivity of these probes, we performed a competitive ABPP-experiment (Figure 1a, b). We incubated the probes in mouse brain lysate and subsequently post-labeled with a color-complementary broad spectrum fluorophosphonate (FP) probe before visualizing the targets on SDS-PAGE.^12^ This allowed simultaneous monitoring of MAGL-labeling and inhibition. Direct fluorescent read-out of the SDS-PAGE showed that MAGL was efficiently targeted at concentrations as low as 10 nM (Figure 1c). Furthermore, the probes outcompeted FP-probes with a IC_50_ of ∼100 nM. At high concentrations an off-target at ∼70 kDa was observed, which is likely to be one or more of the carboxylesterases, a common off-target of activated carbamates.^38^

We further assessed the selectivity of the probes by incubating the probes in brain lysates from wild-type or *Mgll*-/- mice (Figure 1d). No labeling was observed in lysates from *Mgll*-/- mouse brains, indicating selective labeling of MAGL at 100 nM (Figure 1d). Further profiling in both mouse and human brain lysates (Figure 1e-f) showed selective targeting of MAGL in both species, where the labeling could be ablated by selective MAGL inhibitors (ABX-1431, LEI-515) and the non-fluorescent probe **1**, but not by inhibitors of the other 2-AG hydrolases ABHD6 (KT182) or ABHD12 (DO264). In conclusion, the developed probes label active MAGL in mouse and human brain lysates with high selectivity.

### Cellular labeling of monoacylglycerol lipase

After validating this set of active fluorescent probes *in vitro*, their performance in living cells was investigated. The human neuroglioma cell line U-87 MG was chosen for probes targeting MAGL, as these cells have high MAGL expression.^42^ U-87 MG cells were incubated with increasing concentration of fluorescent MAGL-probes, followed by lysis and analysis of the lysates by SDS-PAGE and in-gel fluorescent scanning.

This revealed effective intracellular labeling of MAGL by all three probes at concentrations as low as 1 nM (Figure 2a). Apparent EC_50_ values were around 10 nM, with **LEI-463-BDP-TMR** and **LEI-463-BDP-FL** showing slightly higher potency than **LEI-463-Cy5** (Figure 2b). To capture both inhibitory- and activating effects on MAGL activity in this cell line we chose 10 nM probe (∼50% target engagement) as the optimal concentration. Incubating cells with this concentration produced a strong signal on SDS-PAGE after in-gel fluorescent scanning, which was completely ablated by pre-incubation with the covalent MAGL inhibitor ABX-1431 (Figure 3c). Dose-response incubation with ABX-1431 demonstrated target engagement with a pIC_50_ value of 8.6 ± 0.2 (Figure 2d, e). These results indicate that competitive ABPP using **LEI-463** probes is an effective method for evaluating the potency of MAGL inhibitors on endogenous MAGL in living cells. Recently, an *in situ* target engagement assay for MAGL was described by Gazzi *et al*.^43^ This assay, while allowing for a high throughput, requires ectopic expression of a MAGL-Nanoluciferase fusion construct, while our method works on endogenous MAGL.

**Figure 2.**
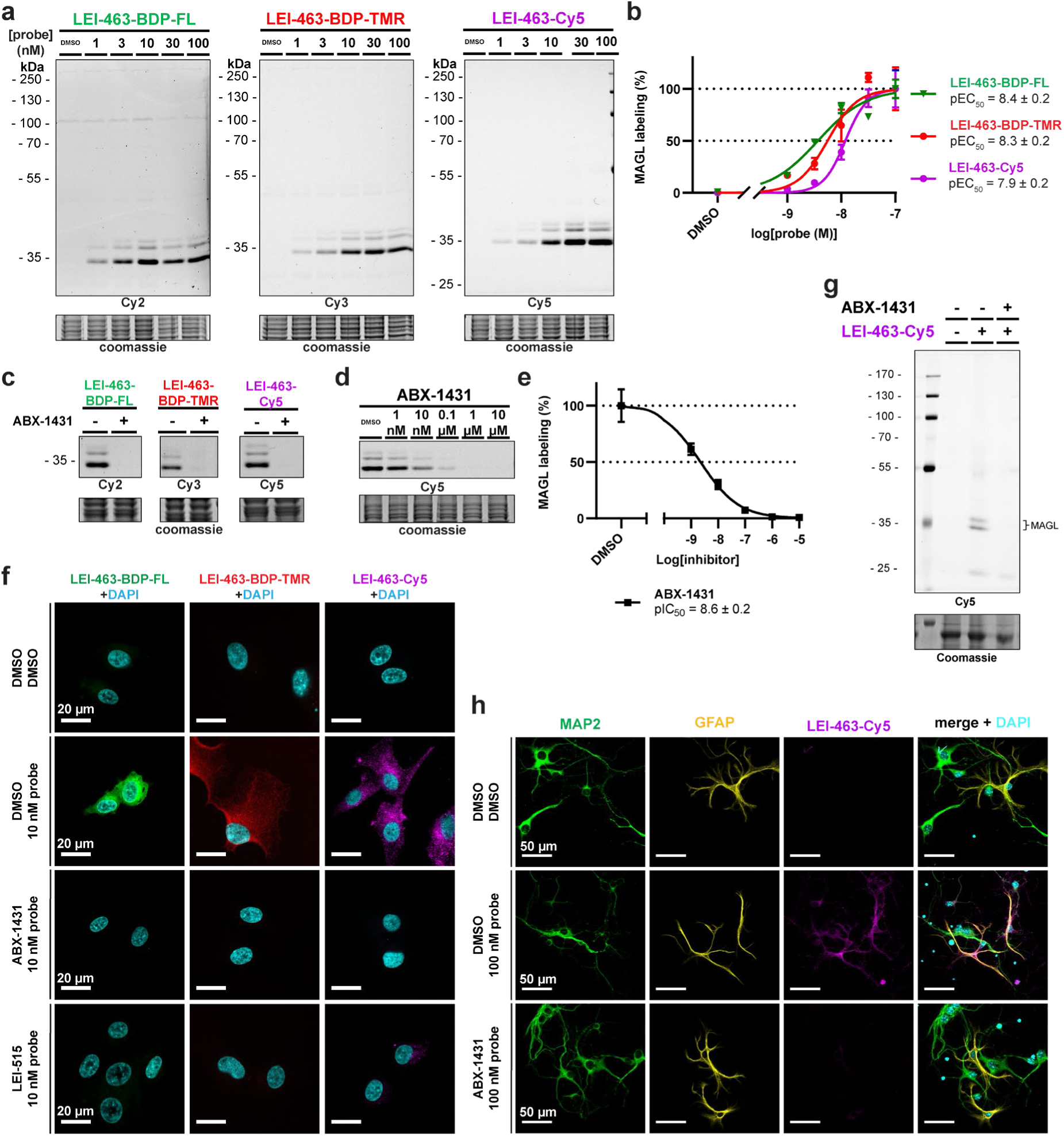
Targeting MAGL in living U-87 MG cells and primary cortical cultures. **a**, Dose-response engagement of MAGL in U-87 MG cells. Cells were incubated with the indicated probes for 1 hour at 37 °C. **b**, Quantification of the fluorescent signal with the signal at 100 nM set to 100% (n=3). **c,** Fluorescent MAGL bands were outcompeted with ABX-1431 (10 µM, 1h, 37 °C). **d,** Dose-response inhibition of MAGL *in situ*by ABX-1431. Cells were incubated with the indicated doses of ABX-1431 for 1 hour before treatment with probe (1 hour, 10 nM, 37 °C). **e**, Quantification of **d** (N=3). **f,** Cells were incubated with MAGL probes (10 nM, 1 hour, 37 °C) and subsequently fixed, stained for DAPI (nuclei, cyan) and imaged with confocal fluorescent microscopy. Scale bars are 20 µm. **g-h**, Primary cortical neuron-astrocyte co-cultures were treated with DMSO or ABX-1431 (10 µM, 1h) before targeting MAGL with **LEI-463-Cy5** (100 nM, 1h, 37 °C). The cultures were then lysed and analyzed on SDS-PAGE (**g**) or fixed, stained for GFAP (astrocytes, yellow), MAP2 (neurons, green) and DAPI (nuclei, cyan) and imaged by confocal microscopy (**h**). Scale bars are 50 µm.

**Figure 3.**
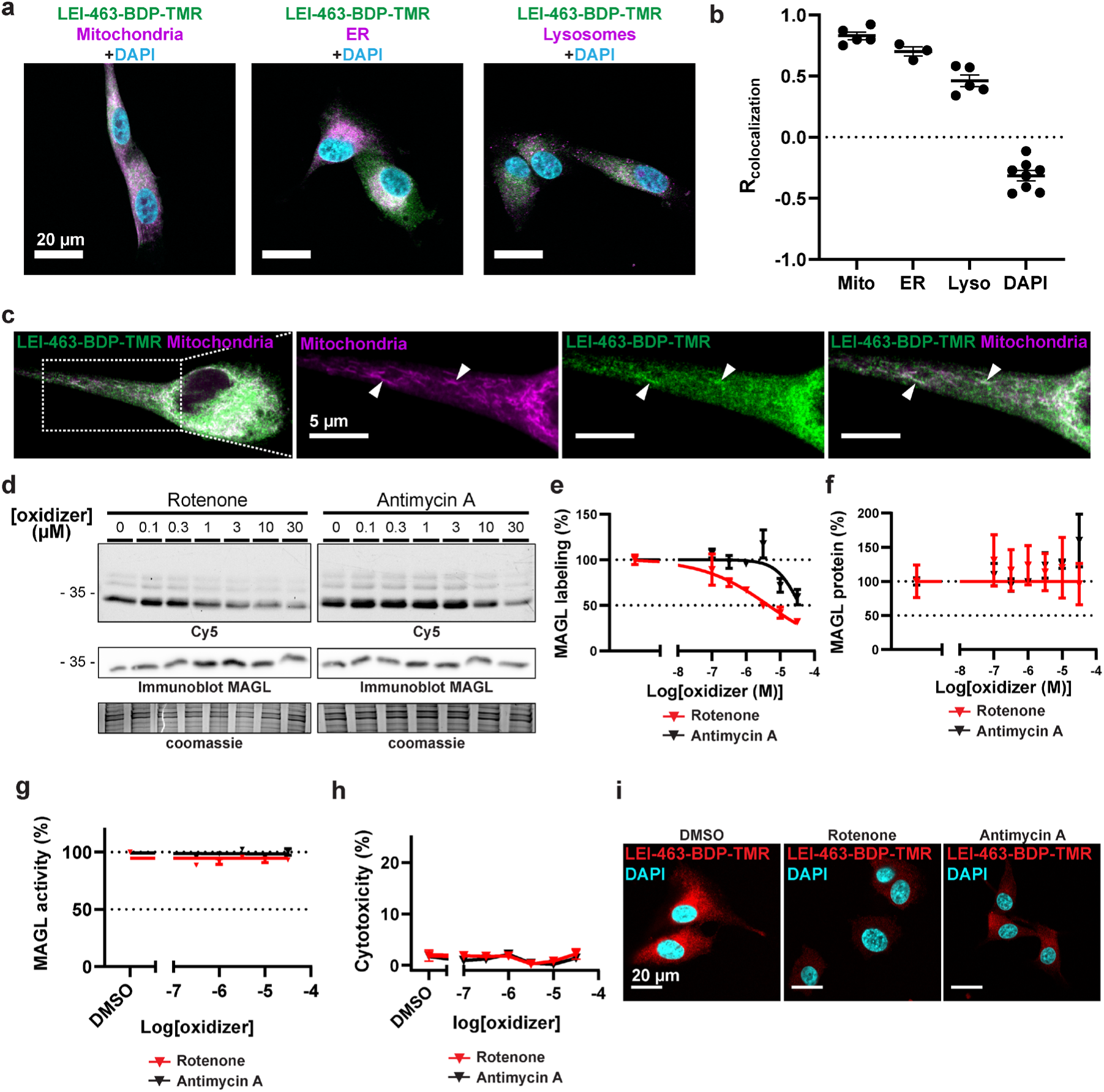
MAGL activity is affected by mitochondrial disruption. **a,** U-87 MG cells were incubated with **LEI-463-BDP-TMR** and combined with a staining for organelles. Scale bars are 20 µm. **b,** Quantification of colocalization as measured by Pearson’s coefficient. One data point is one measured cell (n=3-8 cells). **c,** Zoom of MAGL-mitochondria colocalization. Scale bars are 5 µm. **d,** U-87 MG cells were treated with increasing doses of rotenone or antimycin A for 2 hours and subsequently treated with 10 nM **LEI-463-Cy5** to measure levels of active MAGL. **e,** Quantification of **LEI-463-Cy5** signal from **d** (n=3). **f**, Quantification of immunoblot signal from **d** (n=3). **g,** *In vitro* activity of MAGL is not affected by rotenone and antimycin A. MAGL activity was measured in membranes from lysed MAGL-overexpressing HEK293T cells using a natural substrate assay^42,44^ (N=2, n=2). **h,** Viability of U-87 MG cells after treatment with oxidizers (n=3). **i,** Cells were treated with antimycin A or rotenone (2h, 10 µM) and subsequently treated with **LEI-463-BDP-TMR**. MAGL activity was imaged using a confocal microscope. Scale bars are 20 µm.

Next, the fluorescent signal was visualized using confocal microscopy (Figure 2f). In U-87 MG cells, intracellular signal showed prominent labeling in a cytosolic punctate pattern. This fluorescent signal was inhibited by MAGL inhibitors **ABX-1431** and **LEI-515**, confirming its specificity. Lastly, the ability of **LEI-463** probes to target MAGL in a more complex and physiological system was investigated. Primary cultures of neonatal mouse cortex, containing a mixture of cortical neurons and astrocytes, were prepared. Treatment of these cultures with **LEI-463-Cy5** resulted in selective engagement of MAGL as analyzed by SDS-PAGE and in-gel fluorescent scanning (Figure 2g) and fluorescence microscopy (Figure 2h). Notably, glial fibrillary acidic protein (GFAP) positive astrocytes exhibited high levels of MAGL activity in these cultures (Figure 2h).

To investigate the identity of intracellular organelles on which active MAGL resides, U-87 MG cells were treated with 10 nM **LEI-463-BDP-TMR** and co-incubated with fluorescent trackers for lysosomes, the endoplasmic reticulum (ER) and mitochondria (Figure 3a). This experiment revealed partial co-localization with both mitochondria and the ER (Figure 3a-c). However, much of the intracellular signal accumulated in a punctate pattern that did not colocalize with any of these organelles. It is speculated that these might be lipid droplets, as MAGL in adipocytes is involved in neutral lipid cycling on lipid droplets and these are organelles with high monoacylglycerol levels.^45^

Dotsey *et al.* have previously demonstrated that oxidative stress in cells can lead to oxidation of two regulatory cysteines in the lid-domain of MAGL, thereby disrupting its activity.^46^ To explore whether this post-translational modification of MAGL activity can be detected by our probe, we incubated a small set of oxidative stress inducers in the U-87 MG cells. Rotenone and antimycin A - both blockers of mitochondrial respiration and strong inducers of ROS^47^ - reduced MAGL labeling by **LEI-463-Cy5** (Figure S1) in a concentration-dependent manner (Figure 3d, e), whereas MAGL protein levels remained unchanged as quantified by immunoblots against MAGL (Figure 3d, f). Rotenone and antimycin A did not act as direct inhibitors of MAGL (Figure 3g) and no cytotoxicity was observed (Figure 3h). Furthermore, no change in the subcellular distribution of MAGL was noted (Figure 3i). Taken together, these results suggest that MAGL activity is inhibited by rotenone and antimycin A through a post-translational mechanism, most likely involving oxidative stress that oxidizes regulatory cysteines in the lid-domain of MAGL.^46^ These findings highlight the ability of the **LEI-463** probes to monitor MAGL activity in cellular processes, complementing classical techniques that measure only protein expression levels.

### Activity-based histology of MAGL in the mouse brain

We next sought to develop a protocol to image MAGL activity in tissues, here termed *activity-based histology*. Initial experiments to develop an activity-based histology protocol focused on fresh-frozen mouse brain tissue. Unfixed parasagittal cryosections from fresh-frozen mouse brains were incubated with MAGL probe **LEI-463-BDP-TMR**. To control for unspecific signal, sections were pre-incubated with ABX-1431. However, unspecific labeling occurred in ABX-1431 treated sections when the tissue was only washed with aqueous buffers like PBS, even when supplied with the detergent Triton-X100 (Figure S2). Given the high lipophilicity of the **LEI-463** probes (LogP 7-8) this most likely occurred because the probe has unspecific lipophilic interactions with the lipid-rich environment of the brain, most notably myelin. In line with this premise, washing the tissue in 100% methanol, which dissolves the lipophilic probes, did remove unspecific signal, while preserving signal in granule cell layer and the molecular layer of the cerebellum (Figure S2).

Although washing the tissue in methanol allowed visualization of specific signal, tissue integrity was compromised, limiting high-resolution imaging. Surprisingly, a short fixation with 4% paraformaldehyde (PFA) prior to probe labeling did not ablate probe labeling. Conversely, this fixation step preserved tissue integrity (Figure S3). In addition, the recently reported EZ Clear protocol^48^, which entails washing sections with a 1:1 solution of tetrahydrofuran (THF) and water, was considered as an alternative for washing tissues in methanol. This method yielded specific probe signal even with high probe concentrations (Figure S4). In contrast to methanol washing, this method does not dehydrate the tissue, preserving integrity and antigen immunogenicity.^48^ Using this optimized protocol (Figure 4a), MAGL activity was visualized in the mouse hippocampus with all three **LEI-463** probes (Figure 4b). Importantly, this signal was absent in the hippocampus of mutant mice lacking MAGL (*Mgll*-/- mice) or in sections treated with ABX-1431 before probe labeling (Figure S5). MAGL activity could also be imaged in other brain regions, such as cerebral cortex, cerebellum and spinal cord (Figure S6).

**Figure 4.**
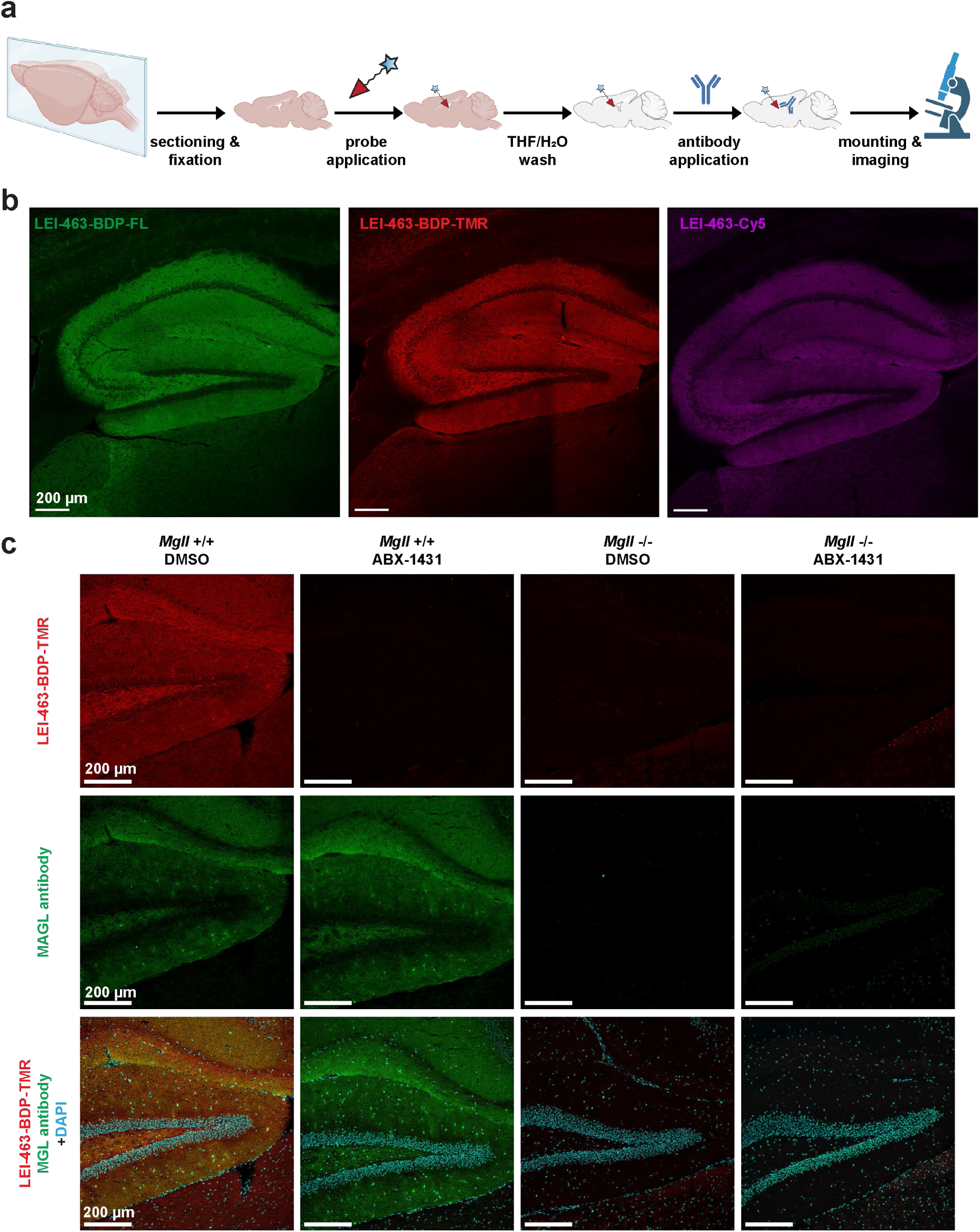
Activity-based histology of MAGL in the mouse brain. **a**, Workflow of the activity-based histology method. **b**, Fresh-frozen cryosections from mouse brain were treated with the indicated **LEI-463** probe (1 µM, 30 min, RT) and the hippocampus was imaged with confocal microscopy. **c**, **LEI-463-BDP-TMR** staining was combined with an antibody staining against MAGL in WT and MAGL knock-out mice (*Mgll-*/-), with and without ABX-1431 treatment, and the dentate gyrus was imaged with confocal microscopy. DAPI shows nuclei. Representative images are shown from 4 independent wild-type and *Mgll*-/- pairs of mice. All scale bars are 200 µm.

Subsequently, probe labeling was combined with immunolabeling against MAGL using an affinity-purified polyclonal antibody^49^. The distribution of MAGL was similar with both staining techniques, showing strong signal in the neuropil of the molecular layer and the hilus of the dentate gyrus. Pre-incubation with ABX-1431 abolished the probe signal, while the MAGL antibody signal persisted. In *Mgll*-/- mice both the probe and immunostaining signals were absent (Figure 4c). This shows that the antibody targets MAGL in both active and inactive states, while the probe only targets active MAGL.

### Distribution of MAGL activity in the mouse hippocampus

With the specificity of the activity-based histology procedure established, **LEI-463**-Cy5 was used to localize active MAGL in specific cell types and neuronal compartments. Previous work by Uchigashima *et al*. utilized traditional immunohistochemical methods in the mouse hippocampus to co-localize MAGL with various cellular and neuronal markers, finding high MAGL protein levels in astrocytes and certain synapse types^49^. Consistent with these findings, the probe labeled numerous astrocyte-like cells in the molecular layer of the dentate gyrus (Figure 5a). Double staining with **LEI-463-Cy5** and glial fibrillary acidic protein (GFAP), an astrocyte marker, confirmed high MAGL activity in astrocytes (Figure 5b).

**Figure 5.**
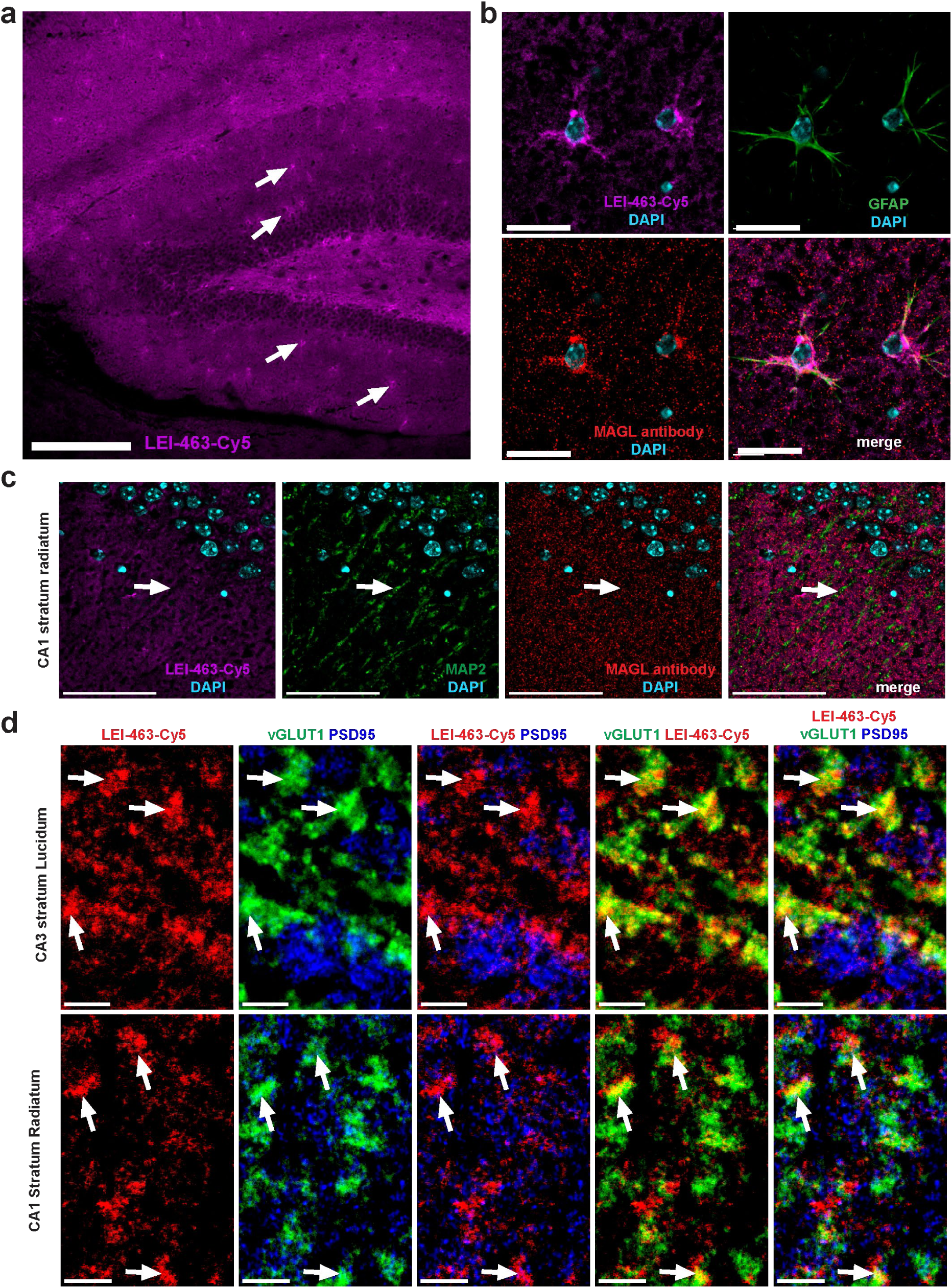
Exploration of the anatomy of MAGL activity throughout the mouse hippocampus. **a**, Overview of **LEI-463**-Cy5 staining in the hippocampal dentate gyrus. **b**, High resolution imaging of MAGL expressing astrocytes in the molecular layer of the dentate gyrus using **LEI-463**-Cy5, a MAGL antibody and GFAP. **c**, Staining with **LEI-463**-Cy5, combined with antibodies against MAP2 and MAGL in the stratum radiatum of the hippocampus. scale bars, 200 µm, **d**, Staining with **LEI-463**-Cy5, combined with antibodies against vGLUT1 and PSD95 in the stratum lucidum of the CA3 (upper panels) or the stratum radiatum of CA1 (lower panels). Scale bars are 200 µm in **a**, 20 µm in **b,** 50 µm in **c,** and 5 µm in **d**.

No co-localization of MAGL activity was found with microtubule-associated protein 2 (MAP2), a marker for neuronal dendrites, in the CA1 region of the hippocampus (Figure 5c), which was in line with previous studies indicating that MAGL predominantly resides in the axonal and pre-synaptic compartment of neurons.^49–52^ A closer examination of MAGL localization at pre- and postsynaptic regions was conducted by combing **LEI-463-Cy5** staining with antibodies for post-synaptic density protein 95 (PSD95), and vesicular glutamate transporter 1 (vGLUT1), markers for post-synaptic regions and excitatory axon terminals, respectively. Significant co-localization between MAGL activity and vGLUT1, but not with PSD95, was found in the stratum lucidum of the CA3 region (Figure 5d). Similarly, the overlap between MAGL activity and vGLUT1-positive Schaffer collateral terminals was observed in the CA1, albeit less than in the stratum lucidum of the CA3 (Figure 5d).

Of note, at higher magnification differences were noted in the staining pattern of the MAGL probe and the antibody, revealing the probe to have a more uniform signal, while antibodies displayed a more punctuated distribution (Figure 5b, c). In conclusion, activity-based histology could visualize MAGL activity distribution in the mouse hippocampus with high spatial resolution. The results were in line with previous reports using traditional immunohistology methods^49,52,53^, showing that MAGL is present and active in hippocampal astrocytes and presynaptic compartments.

### Activity-based histology of MAGL in the human cerebral cortex

Next, the aim was to conduct activity-based histology in human brain samples. Cryosections, obtained from the cortical superior parietal gyrus of donors without neurological disease, were treated with **LEI-463-Cy5** following the same protocol used for mouse tissue (Figure 4a). While fluorescent probe signal was detected in the grey matter, strong, non-specific punctuated signal was still observed in sections treated with ABX-1431 (Figure 6a). This non-specific signal appeared across all wavelengths (Figure S7-S9) and even in untreated sections (Figure S10). This suggests that this signal originates from lipofuscin autofluorescence, a protein that accumulates in cytoplasmic puncta in aging brains.^54^ This type of autofluorescence can be quenched with lipid dyes such as Sudan Black B (SBB) and TrueBlack (TB)^54^, but this did not enable specific detection of **LEI-463** signal, despite effective lipofuscin suppression (Figures S7-S9).

**Figure 6.**
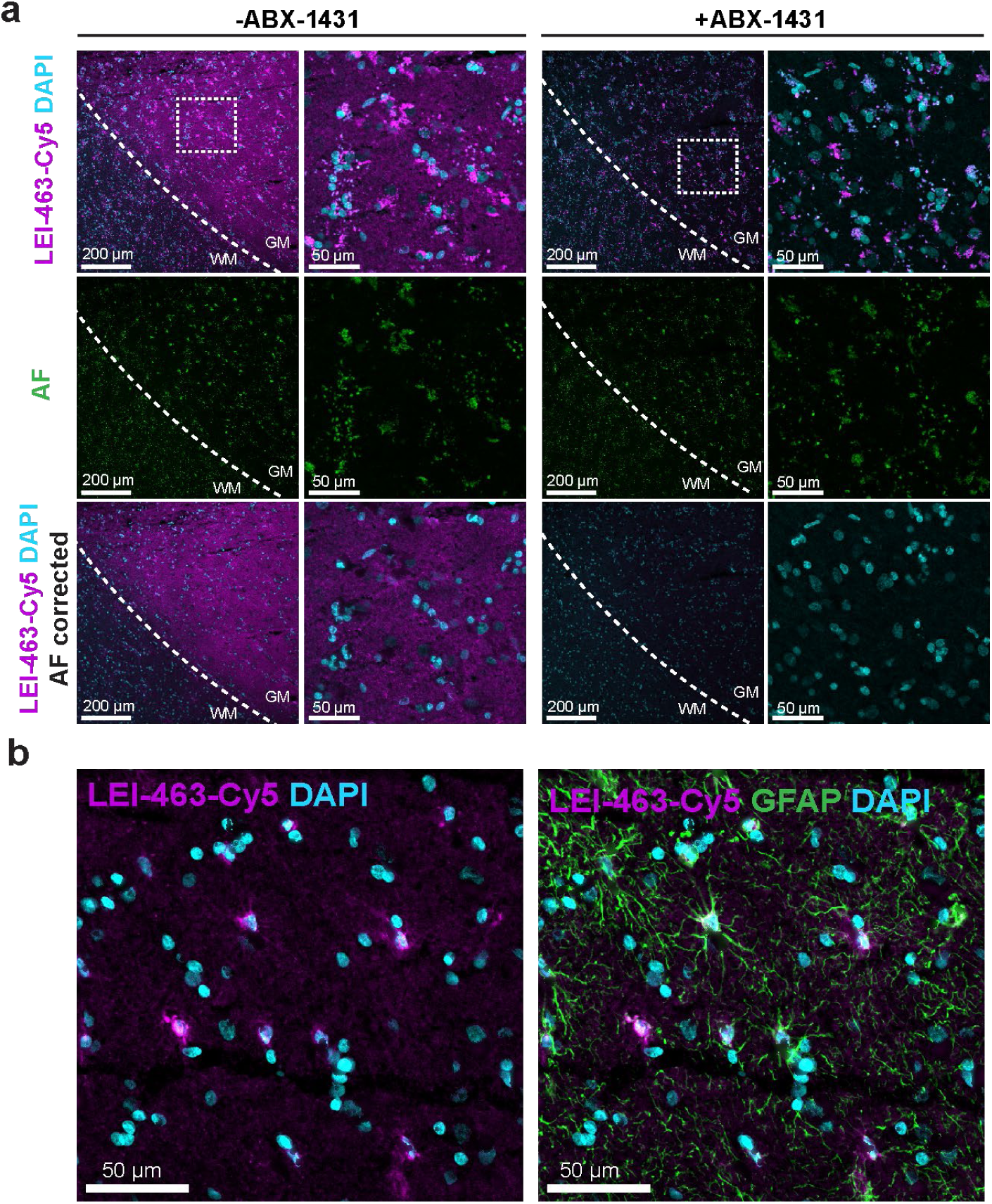
Activity-based histology of MAGL in the human cerebral cortex. **a**, computational autofluorescence removal yields specific MAGL signal in the human cerebral cortex. Sections were treated with **LEI-463-Cy5** as described for the mouse brain, with and without ABX-1431 pretreatment. Autofluorescence (AF) was imaged in the Cy2 channel, which was used to correct the autofluorescence observed in the Cy5 channel (lower panels). The box indicates the region of the higher magnification image. WM indicated the white matter area, while GM indicates cortical grey matter. A representative set of images is shown from 4 independent sets of images. **b**, Co-localization of **LEI-463-Cy5** and GFAP. GFAP was imaged in the Cy2 channel, Autofluorescence was captured in the Cy3 channel (not shown) and used to corrected AF in the other channels.

Consequently, autofluorescence was computationally corrected using signal from other wavelengths. Of note, the autofluorescence in the wavelengths corresponding to the BODIPY-based probes was too intense relative to probe signal, precluding visualization, but this correction enabled the selective visualization of MAGL activity in cortical grey matter with **LEI-463-Cy5** (Figure 6a). Similar results were found across four independent brain donors (Figures S10). Consistent with results in mouse hippocampus, **LEI-463-Cy5** colocalized with GFAP, showing that active MAGL was densest in astrocytes within the human cortex (Figure 6b).

## Discussion

The ability to visualize MAGL activity across the brain at high spatial resolution represents a significant advancement for translational research, particularly in the context of developing MAGL inhibitors as therapeutics. Several such inhibitors are currently undergoing clinical evaluation for a range of neurological and inflammatory conditions. To inform and support these clinical efforts, it is essential to characterize the anatomical, cellular, and subcellular distribution of MAGL activity in both healthy and diseased brain tissue.. In this study, we demonstrate that selective fluorescent activity-based probes are powerful tools to for mapping MAGL activity with high spatial resolution. The **LEI-463** probe series effectively targeted MAGL *in vitro,* in immortalized cell lines and primary cortical cultures. Importantly, we investigated post-translational regulation of MAGL, particularly the redox-sensitive cysteines identified by Dotsey *et al*.^46,55^ Exposure to mitochondrial disruptors such as rotenone and antimycin A, which are known inducers of mitochondrial superoxide production^47^, reduced probe labeling without altering MAGL protein levels, suggesting redox-dependent inhibition. This supports the utility of activity-based probes in capturing dynamic regulatory states of enzymes that would not be evident from expression data alone.^56^

We next developed an *activity-based histology* protocol that enabled visualization of MAGL activity in cryosections from fresh-frozen mouse and human brain tissues. This approach enabled detailed mapping of MAGL activity in the mouse hippocampus, where we observed prominent labeling in astrocytes and presynaptic compartments. Notably, to our knowledge, this is the first report of ABP-based imaging of enzyme activity in human brain tissue. This methodological advancement opens the door to studying MAGL activity distribution in postmortem material from patients with neuroinflammatory and neurodegenerative diseases, including Alzheimer’s disease and multiple sclerosis.

A key limitation of this activity-based histology method is the relatively low signal intensity of **LEI-463** probes compared to traditional antibody-based immunolabeling. Conventional immunostaining benefits from high signal amplification through multiple secondary antibodies binding to each primary antibody, each carrying multiple fluorophores. In contrast, **LEI-463** probes label MAGL with a single fluorophore per active enzyme molecule, potentially explaining the less punctuated staining pattern observed with the probe compared to the antibody (Figure 3c, 4a). Antibodies might produce a more punctate pattern due to sparse labeling events, possibly due to steric hindrance of large antibodies at epitopes, but high signal amplification at each binding event.

Another challenge is the interference from tissue autofluorescence—especially lipofuscin—in aged human brain samples. Although computational methods for autofluorescence subtraction were partially effective, they present limitations: (1) overlapping signal and autofluorescence can lead to data loss; (2) one imaging channel must be dedicated to autofluorescence capture, limiting multiplexing; and (3) regions with low MAGL expression may be obscured. Future improvements should focus on chemical quenching strategies that suppress lipofuscin autofluorescence without compromising probe signal and probe optimization.

Applying this protocol to fresh-frozen cryosections facilitates the use of archived mouse and human tissue samples, which are frequently preserved in this format in brain banks.. Unlike the recent CATCH method by Pang *et al*.^17^, our approach does not require live animal administration or complex tissue clearing, making it particularly suited for retrospective studies on human disease material. Moreover, the compatibility of LEI-463 probes with short formaldehyde fixation suggests potential for extension to perfused tissues, improving morphological preservation without loss of enzymatic signal.

In conclusion, we demonstrate that selective activity-based probes can enable histological mapping of MAGL activity in the mammalian brain with high spatial precision. These tools hold promise for investigating MAGL function in disease models and human pathology, ultimately contributing to therapeutic development in neurodegenerative and neuroinflammatory disorders. More broadly, this work illustrates the value of chemical probes for enhancing histological techniques and bridging the gap between chemical biology and systems neuroscience.

## Acknowledgements

We are grateful to the brain donors and their families for their commitment to the Netherlands Brain Bank donor program. We thank Prof. dr. Sakimura (Niigata University) for providing the MAGL KO animals and E. Tischler, B. Pintér for their technical support. Research reported in the publication was supported by Oncode Accelerator, a Dutch National Growth Fund project under grant number NGFPO2201. DV and MS acknowledge funding from the Institute of Chemical Immunology (project ICI0000030). IK holds the Naus Family Chair in Addiction Sciences in the Department of Psychological and Brain Sciences at Indiana University Bloomington. This work was funded by the NKFIH EXCELLENCE program 151377 (IK) and the National Institutes of Health grant P30DA056410 (IK) and the Intramural Program of the NIH/NIAAA to PP.

## Methods

### Animals, tissues & chemicals

Animal procedures were approved by the Ethics committee for Animal Experiments and the Animal Welfare Body of Leiden University (AVD10600202215851; 15851,1-193) and were performed in accordance with the guidelines of the Dutch government and the European Directive 2010/63/EU. Wild-type C57BL/6J mice were bred in-house and kept in a temperature-controlled room with a 12 hours-12 hours light-dark cycle. Food and water were provided ad libitum. Wild-type brains were obtained from 27 week old C57BL/6 mice, which were killed through CO_2_ inhalation, and subsequently brains and spinal cords were removed and quickly frozen with liquid nitrogen. Subsequently, brains were stored at -80 °C.

MAGL knock-out brains from 10-13 week old mice, alongside wild type littermate control brains, were obtained from Hungary^54^. The mouse line was a kind gift of Kenji Sakimura (Niigata University). Generation of this line has been described in detail previously (Uchigasima 2011). To obtain tissue samples, KO and WT mice were euthanized by decapitation under deep isoflurane anesthesia. Brains were carefully extracted and rapidly frozen with liquid nitrogen. Animal experiments carried out in Hungary were approved by the Hungarian Committee of the Scientific Ethics of Animal Research (license number: MÁB-2018/1) and were conducted according to the Hungarian Act of Animal Care and Experimentation (1998, XXVIII, Section 243/1998, renewed in 40/2013). These comply with the European Communities Council Directive of November 24, 1986 (86/609/EEC; Section 243/1998).

Human brain samples, specifically from the superior parietal gyrus, from four non-demented control donors were obtained from the Netherlands Brain Bank (NBB, www.brainbank.nl). Fresh frozen tissue blocks were dissected, snap-frozen in liquid nitrogen and stored at -80 °C. All procedures of the NBB were approved by the ethical committee of Medical Ethical Committee of Amsterdam Academic Medical Centre and all donors gave written consent to the NBB for the use of their data and tissue for research. Details on the four donors can be found in table 1.

**Table 1.** Donor information.

| Donor No. | Sex (M/F) | Age (years) | PMD (h:min) | pH CSF | Brain weight (g) |
| --- | --- | --- | --- | --- | --- |
| Donor 1 | M | 96 | 06:30 | 6.65 | 1300 |
| Donor 2 | F | 64 | 05:40 | 6.35 | 1221 |
| Donor 3 | M | 102 | 05:00 | 6.64 | 1124 |
| Donor 4 | F | 57 | 07:40 | 6.47 | 1339 |

*PMD: post-mortem delay*

The synthesis and characterization of all activity-based probes used herein is fully described in the supplementary information based on previously published methods^2,38,58,59^. BODIPY-TMR azide and - alkyne were purchased from Lumiprobe (cat. 32430 and cat. D24B0, respectively). Cy5-alkyne were synthesized from the free carboxylic acid, which was provided kindly by Dr. Marta Artola (Leiden University). FP-BODIPY, FP-TAMRA, MB064, ABX-1431 and LEI-515 were synthesized in house as previously described^12,37,42,60^. KT182 was purchased from Merck (SML1248) and DO264 was purchased from Focus Biomolecules (10-3712). Buthionine Sulfoximine (BSO) was used as purchased from Cayman Chemicals. H_2_O_2_, carbonyl cyanide m-chlorophenyl hydrazone (CCCP), Antimycin A, 1-Methyl-4-phenyl-1,2,3,6-tetrahydropyridine (MPTP) hydrochloride, and rotenone were purchased from Sigma Aldrich. H_2_O_2_ was kept as a 9.8 M stock in Milli-Q demineralized water. MPTP was dissolved in Milli-Q demineralized water as a 10 mM stock. BSO was dissolved in Milli-Q demineralized water as a 100 mM stock. CCCP, rotenone, and antimycin A were dissolved in dimethylsulfoxide (DMSO) as 10 mM stocks. Used organelle stainings were LysoTracker™ Green DND-26 (ThermoFisher) 1mM in DMSO (∼504/511 nm), ER-Tracker™ Green BODIPY™ FL Glibenclamide (ThermoFisher) (∼504/511 nm) and MitoTracker™ Deep Red FM (ThermoFisher) (∼644/665 nm). The used antibodies are listed in Table 2.

**Table 2.**
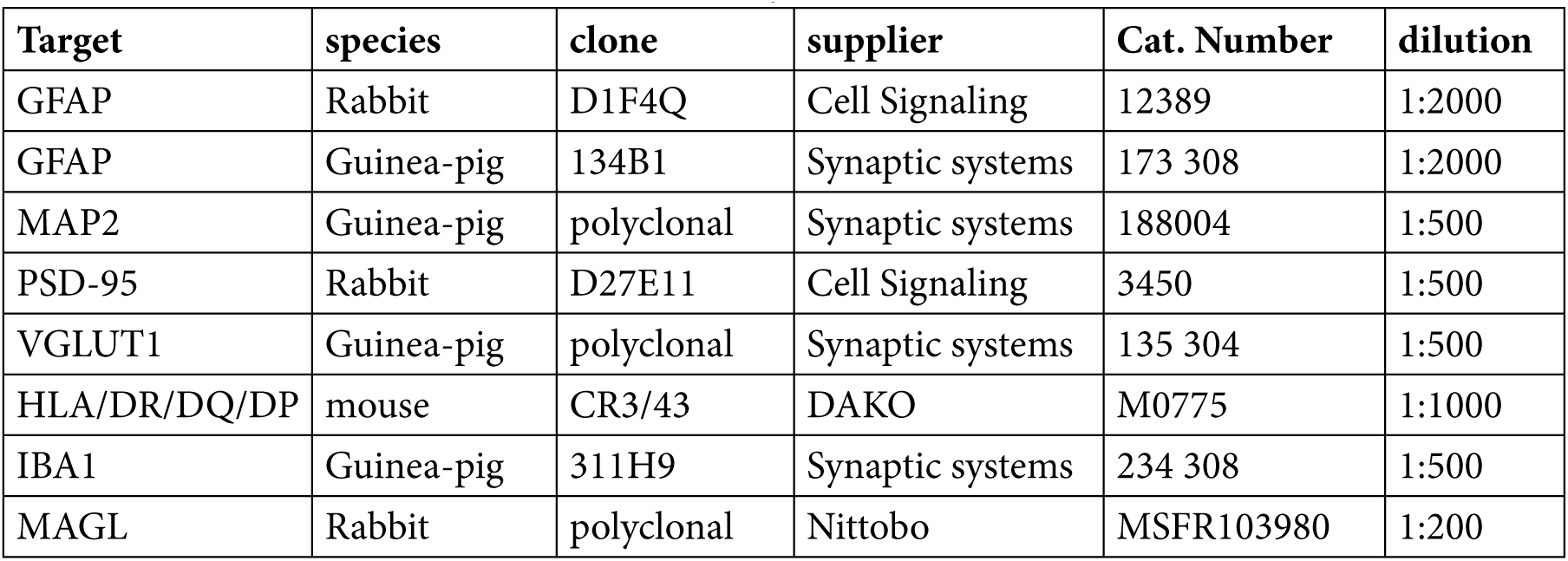
Overview of used antibodies in this study.

| Target | species | clone | supplier | Cat. Number | dilution |
| --- | --- | --- | --- | --- | --- |
| GFAP | Rabbit | D1F4Q | Cell Signaling | 12389 | 1:2000 |
| GFAP | Guinea-pig | 134B1 | Synaptic systems | 173 308 | 1:2000 |
| MAP2 | Guinea-pig | polyclonal | Synaptic systems | 188004 | 1:500 |
| PSD-95 | Rabbit | D27E11 | Cell Signaling | 3450 | 1:500 |
| VGLUT1 | Guinea-pig | polyclonal | Synaptic systems | 135 304 | 1:500 |
| HLA/DR/DQ/DP | mouse | CR3/43 | DAKO | M0775 | 1:1000 |
| IBA1 | Guinea-pig | 311H9 | Synaptic systems | 234 308 | 1:500 |
| MAGL | Rabbit | polyclonal | Nittobo | MSFR103980 | 1:200 |

### Tissue lysate preparation

Mouse brains were removed from adult (27 week old) male C57BL/6 mice, who were sacrificed by inhalation of CO_2_. Whole brains were removed, snap-frozen in liquid nitrogen and stored at -80 °C until use. For preparation of lysates, brains were thawed on ice and added was ice-cold lysis buffer (20 mM HEPES, pH 7.2, 1 mM MgCl_2_, 2 U/mL benzonase), 1.0 mm glass beads and the tissue was homogenized in a bullet blender (3x30s). Debris was subsequently removed by short centrifugation at 4 °C (2 min, 3000 rpm). The supernatant was transferred to new tubes, the protein concentrations was determined using a Bradford assay (Bio-Rad) and the lysates were diluted with 20 mM HEPES buffer to 2 mg/mL protein.

To obtain lysates for human cortical tissue, tissue blocks were sectioned at 50 μm in a cryostat (Thermo Fisher HM525 NX) at -20 °C with Thermo Fisher MX35 Ultra microtome blades. To prepare homogenates, sections were weighed and subsequently lysed in ice cold lysis buffer (20 mM HEPES, pH 7.2, 1 mM MgCl2, 2 U/mL benzonase) at 12 μL/mg tissue. Glass beads (1.0 mm, BioSpec, cat. No 11079110) were added and the tissue was homogenized in a bullet blender (3x30s). The lysate was transferred to new Eppendorf tubes and the protein concentrations were measured using a Bradford assay and protein concentrations were adjusted to 1 mg/mL using dilution buffer (20 mM HEPES, pH 7.2, 2 mM DTT, 1 mM MgCl_2_. Lysates were snap frozen in liquid nitrogen and then stored at -80 °C.

### *In vitro* activity-based protein profiling

Proteomes were thawed on ice and brought to RT. 19 µL of lysate was incubated with probe (1 µL from a 20x stock in DMSO) or with inhibitor first (0.5 µL from a 40x stock) and subsequently with probe (0.5 µL from a 40x stock). MAGL-probes (**LEI-463**) were incubated at RT for 30 min. After incubation, the reaction was quenched through the addition of 7.5 µL of 4x Laemmli buffer (final concentrations: 60 mM Tris (pH 6.8), 2% (w/v) SDS, 10% (v/v) glycerol, 1.25% (v/v) β-mercaptoethanol, 0.01% (v/v) bromophenol blue) and the lysate was loaded on 10% AA-gels (29:1 crosslinker) and resolved at 180 V for 70-75 min. The wet gel slabs were subsequently scanned on a Chemidoc MP (Bio-Rad) imaging system on Cy2 (Blue Epi, 532/28 filter, 120s exposure), Cy3 (Green Epi, 602/50 filter, 120 s exposure) or Cy5 (Red Epi, 700/50 filter, 10 s exposure).

### SDS-PAGE image analysis

Images of SDS-PAGE gels were analyzed using Imagelab 6.0 (Bio-Rad). Intensities were corrected for protein loading by coomassie staining and then normalized on the average of the control samples. pEC_50_/pIC_50_ values were calculated using GraphPad Prism 9 using the “variable slope dose-response fit”. For all quantifications at least three replicates of each condition were used. For EC_50_ curves, the highest concentration tested was assumed to be 100%.

### Cell culture and *in situ* treatments

U-87 MG cells were obtained from ATCC and maintained in DMEM high glucose (D6547), supplemented with 10% FCS, P/S and 2 mM Glutamine in a humidified incubator at 37 °C with 7% CO_2_. For passage, cells were washed in PBS, and subsequently detached by trypsinization. After detachment from the culture plate, trypsin was inactivated through the addition of fresh culture medium with serum and cells were passaged as required. All cell cultures were maintained for 2-3 months and then discarded. All cell cultures were regularly tested for mycoplasma infection and consistently tested negative.

U-87 MG cells seeded the day before treatment in a 6-well plate at 2×10^5^ cells in 2 mL culture medium per well, or seeded in a 96-well plate at 3×10^4^ cells in 150 μL culture medium per well. During experiments, cells were kept in DMEM high glucose (Sigma D6547), supplemented with 2% v/v fetal calf serum (FCS), penicillin and streptomycin, 100 μg/mL each, and 2 mM stabilized L-alanyl-L-glutamine from GlutaMAX™ supplement (Gibco) in a humidified incubator at 37 °C with 7% CO2. In 6-well plates, cells were treated in 1 mL treatment medium. In 96-well plates, cells were treated in 100 μL treatment medium. Cells were pre-incubated with medium enriched with H_2_O_2_ and BSO for 30 minutes, ABX-1431, LEI-515 and MPTP for 1 hour, and CCCP, rotenone and antimycin A for 2 hours in a humidified incubator at 37 °C with 7% CO2. Subsequently, treatment medium containing **LEI-463 probe** final concentration (0.1% DMSO) was added and cells were incubated for 1 hour. For the dose-response analysis, probe concentrations are as indicated. For all other analyses, 10 nM of **LEI-463** probe was used.

For gel-based analysis, after the final incubation step, cells were washed with PBS. Successive cell lysis was achieved using SDS lysis buffer (1% v/v SDS, 50 mM Tris and 2 mM EDTA at pH 7) and sonication (25%, 30 seconds) in a Q700 sonicator (QSonica). Protein concentrations were determined using a Pierce protein assay kit (Thermo Fisher) and equal protein loading (8-12 µg protein) were resolved on SDS-PAGE as described above.

For microscopy, sells were seeded in 8-chamber covered μ-slides (80826, IBIDI) at 10^4^ cells in 200 μL culture medium per chamber and subsequently treated as above. For colocalization experiments, cells were incubated with probe for 1 hour, cells were washed with PBS three times, and cells were incubated with 0.1% v/v 1μM LysoTracker™, 1μM ERTracker™ or 25 nM MitoTracker™ according to manufacturer’s recommendations for 30 minutes at 37 °C. Reactions taking place in IBIDI chambers were quenched by replacing medium with PBS. Cells were washed with PBS and fixed in 4% v/v paraformaldehyde in PBS for 15 minutes. Fixed cells were washed with PBS three times, with methanol three times, washed with PBS once and stained using Hoechst 33342 (2 μg/mL in PBS). Cells were finally washed with PBS three times and stored at 4°C in Glycerol-DABCO.

### Western Blot Analysis

After fluorescence scanning of SDS-PAGE gels for gel-based activity-based protein profiling, a selected part of the gel was sliced for Coomassie staining. The remaining gel was placed in a Trans-Blot Turbo Mini 0.2μm PVDF Transfer Pack (Bio-Rad) sandwich and transferred using a Trans-Blot Turbo (Bio-Rad). MAGL was sliced from the membrane and the slice was washed in TBS and TBST (5 min.), 5% milk in TBST for 1 hour, and TBST three times. Cut sections were incubated with rabbit-anti-MAGL (1:200) antibody (ab24701, Abcam) in 5% milk in TBST overnight at 4°C. Membranes were washed with TBST three times, reacted with mouse-anti-rabbit IgG-HRP (1:5000) (Santa Cruz) in 5% milk in TBST for 1 hour, washed with TBST three times, and one final time with TBS. For imaging, membranes were prepared for two minutes in luminol solution, final concentrations 14 mM luminol, 6.7 mM p-coumaric acid and 3 mM H_2_O_2_, and scanned using the Chemidoc MP on the Chemiluminescence channel (646SP, No Light). Quantified Chemiluminescence intensity data was corrected for protein loading by relative Coomassie intensity.

### Cytotoxicity assay

Cytotoxicity of compounds was determined using the CyQUANT™ LDH Cytotoxicity Assay, following the manufacturer’s instructions (ThermoFisher). Cytotoxicity of compounds was determined on fresh cells seeded at 2×10^4^ cells/well by incubating with compounds for 75 minutes, adding medium enriched with lysis buffer to a maximum series and incubating for another 45 minutes. Medium from all samples was incubated with ‘Reaction Mixture’ and reactions were quenched with ‘Stop Solution’. Absorbance measurements were conducted on a CLARIOstar® plate reader (IsoGen).

### Primary cell cultures

Primary neuron/astrocyte cultures were prepared from cortices of 0-2 day old C57BL/6J mice and cultured as described by Straub *et al.*^61^ Cells were seeded at 5×10^5^ cells/well in poly-L-lysine coated 6-well plates (Corning, 3516) and at 3.5×10^4^ cells/well in poly-L-lysine coated μ-slides (80826, IBIDI). Cultures were used at day in vitro 14-15.The cultures were treated either DMSO or ABX-1431 (10 µM) in culture medium for 1 hour, before treatment with 10 nM **LEI-463-Cy5** for an additional hour. Cultures in 6-well plates were analyzed by SDS-PAGE as described for U-87 MG cells. Cultures in μ-slides (80826, IBIDI) were washed with PBS, fixed in 4% PFA in PBS, subsequently washed with 50% tetrahydrofuran in MilliQ before rehydrating on PBS. Then the cultures were blocked using 10% donkey serum in PBS containing 0.5% Triton-X. Cells were incubated with primary antibodies against GFAP (1:1000, D1F4Q, Cell signaling) and MAP2 (polyclonal guinea pig antiserum, anti-MAP2 – 188 004, Synaptic Systems) in PBS with 5% donkey serum and 0.5% triton-X overnight at 4 °C. The next day, sections were washed in PBS (3x5min), and the sections were incubated with appropriate secondary antibodies (Jackson Immuno, Donkey anti-IgG (H+L), 1:1000) in PBS with 0.5% triton-X. Afterwards, cells were washed with PBS once and stained using Hoechst 33342 (2 μg/mL in PBS). Cells were finally washed with PBS three times and stored at 4°C in 80% Glycerol with DABCO.

### Preparation of tissue cryosections

Fresh frozen brain tissue, stored at -80 °C, were slowly warmed up to an appropriate temperature of -16 to -14 °C for ∼ 1 hour). Then, 20 µm sections were prepared using a cryostat (ThermoFisher HM525NX). Sections were mounted on slides (Epredia, J1800AMNZ) and temporarily stored (max ∼1 hour) in a storage box on ice, before being processed further. Generally, sections were always prepared fresh. However, sections have also been prepared that were dried overnight in a box with dehydrating silica beads, sealed in tube foil and stored at -80 °C. After this procedure, the MAGL probe staining still worked in mouse brain tissue (data not shown). This was not tested for human tissue.

### Activity-based histology procedure using MAGL probes

First, a hydrophobic box was drawn around the tissue sections using a PAP pen (Sigma, Z672548). Each tissue section was fixed with 4% paraformaldehyde (PFA) in PBS for 10 minutes at room temperature (RT). The incubation buffer for treatment was 20 mM HEPES buffer pH7.2 with 1% w/v bovine serum albumin (HEPES-BSA buffer). Next, three HEPES-BSA buffers were prepared: one containing 1% v/v DMSO (vehicle), one with 10 μM ABX-1431, and a third containing 1 µM of a MAGL probe. After fixation, sections were washed with 200 µL of the HEPES-BSA buffer, for 3 min. Sections were further incubated with 200 µL of either the 10 µM ABX-1431 or vehicle solution (30 min, RT). Afterwards, the sections were washed with the HEPES-BSA buffer before sections were incubated with 200 µL of 1 µM MAGL probe (30 min, RT). Sections were washed again with HEPES-BSA buffer once and subsequently in PBS for 3x5 minutes, and lastly once in milliQ water. The sections were then submerged in a 1:1 solution of tetrahydrofuran (THF) and milliQ water and gently rocked for 1 hour, to remove excess probe, while being kept dark. Sections were then washed in milliQ water for 4x5 minutes before being submerged in PBS for 1x5 minutes. Nuclei staining was done using 200 µL of Hoechst 33342 (2 µg/mL) or DAPI (1 µg/mL) in PBS before sections were mounted in Mowiol 4-88 mounting medium (100 mg/mL Mowiol 4-88, 25% glycerol, 25 mg/mL DABCO in 0.1 M Tris-HCl pH 8.5) and stored overnight at 4 °C.

### Microscopy

Images were acquired at a Nikon Ti2-E inverted confocal microscope coupled to a Nikon A1R confocal unit, using Galvano scanning. Used objectives were a HP Apo TIRF 100xAC/1.49 numerical aperture (NA), or a Plan Apo λ 20x objective. Used laser lines were DAPI (405 nm), Cy2 (488 nm), Cy3 (561 nm) and Cy5 (647 nm) in combination with a 405/488/561/647 dichroic mirror system and 450/50, 525/50 and 595/5-filter cubes. Generally, the pinhole was set to 1.2 airy units (AU) based on the longest wavelength included in the experiment. Then, the laser strengths and gains were balanced to obtain strong, but not saturated, signal in the positive control samples. Settings were subsequently kept the same for all samples in the experiment. All images were further processed using ImageJ, where linear adjustments were made to min/max intensities to maximize desired contrast, which were always applied the same to all images of that experiment. The gamma was never adjusted.

### Antibody staining

When probe staining was multiplexed with antibody staining, this procedure was performed after the EZ CLEAR-tissue clearing step. First, 200 µL of 10% donkey serum, 0.5% TritonX-100 in PBS was used to block the tissues in a moist chamber for 2 hours at RT. After blocking, the sections were washed in PBS (3x5 min), before primary antibodies were added which were left to incubate overnight at 4 °C (Table 2).

The next day, sections were washed in PBS (3x5min), and the sections were incubated with appropriate secondary antibodies (Jackson Immuno, Donkey anti-IgG (H+L), 1:1000). Afterwards, sections were mounted and imaged as described above.

### Autofluorescence processing with ImageJ

Autofluorescence subtraction was done using macro code in ImageJ, in short: the channel containing solely autofluorescence (AF) signal was used to generate a mask-selection of the autofluorescent pixels.

The mask was then overlapped with the probe/antibody containing channels. The mean pixel intensity is measured between the inside and outside of the mask, calculating a ratio < 1. The ratio was then applied to correct the pixel intensity inside the AF-mask, by reducing it a similar value as its surroundings. Lastly, a Gaussian Blur was applied of either 2 (zoomed-out images, 20x lens) or 8 (zoomed-in images, 100x lens).

### Statistical methods

Statistical analyses were performed using GraphPad Prism 9.0. Data is always expressed as mean ± SEM. The statistical test used in each experiment is indicated in its respective figure caption. No methods have been employed to determine sample sizes.

