## Supplementary information for "Spatially resolved mapping of monoacylglycerol lipase activity in the brain"

to

**Supplementary figures**

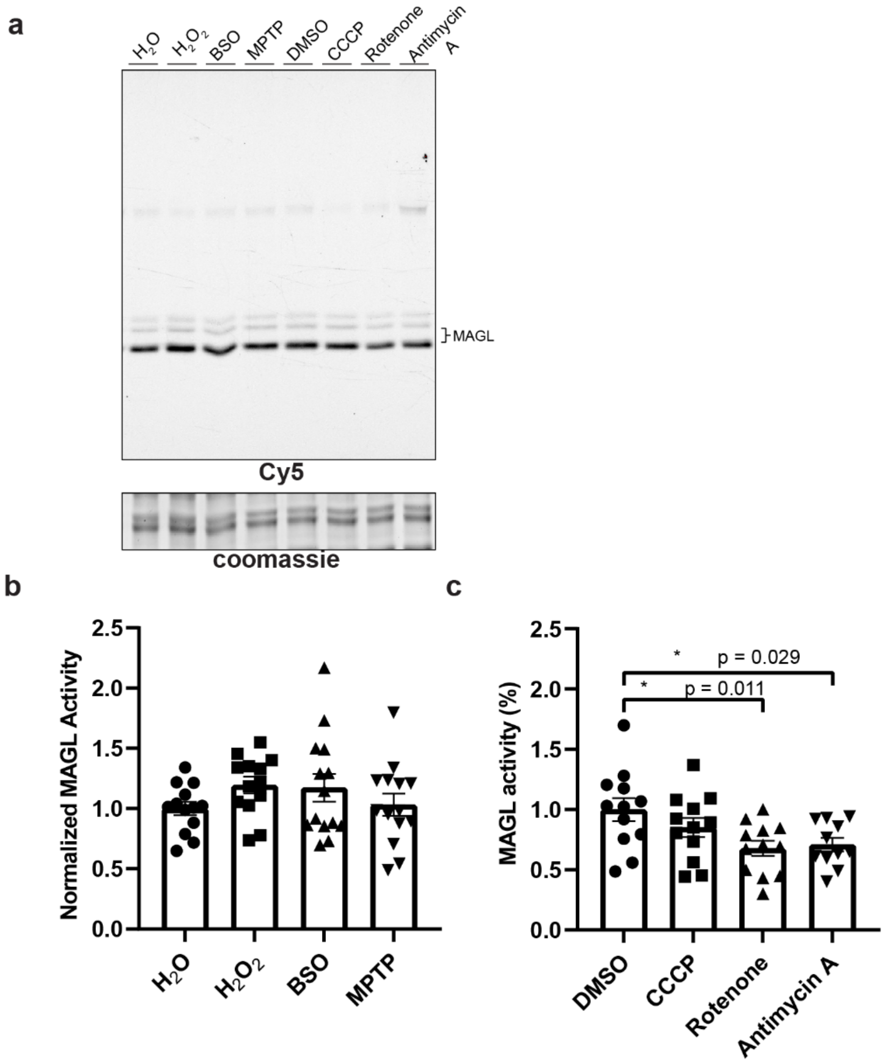


**Figure S1. Oxidizer screening on MAGL activity.** U-87 MG cells were incubated with oxidizing compounds, treated with **LEI-463-Cy5** (10 nM, 1h *in situ*) and resolved on SDS-PAGE. One-way ANOVA with post-hoc Tukey test and Dunett's multiple testing correction was applied.

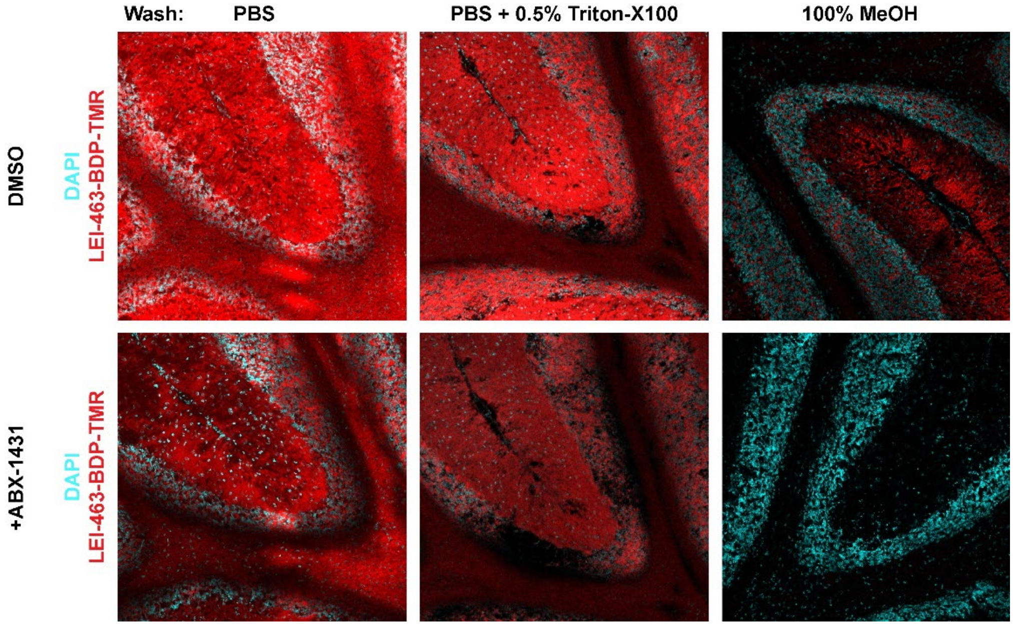


**Figure S2. Unspecific probe signal is retained after aqueous washing.** Fresh-frozen cryosections from mouse brain were treated with either DMSO or ABX-1431 (10 µM, 30min, RT) with LEI-463-BDP-TMR (100 nM, 30 min, RT) and subsequently fixed, washed with the indicated solutions and the cerebellum was imaged with confocal microscopy. Scale bars are 200 µm.

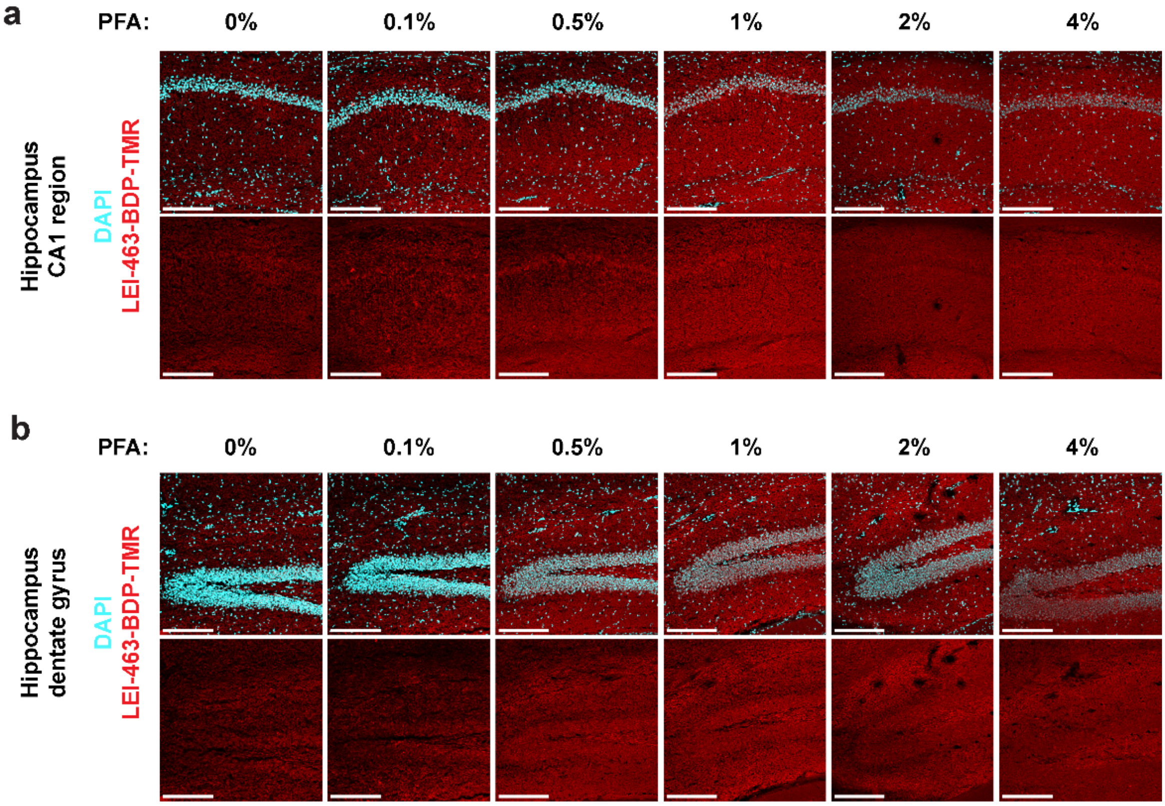


**Figure S3. Formaldehyde fixation does not block probe signal, while improving tissue integrity.** Fresh-frozen cryosections from mouse brain were pre-fixed with the indicated concentrations of PFA in PBS, then treated with LEI-463-BDP-TMR (100 nM, 30 min, RT) and subsequently post-fixed in 4% PFA, washed with methanol and the hippocampal CA1 (**a**) and dentate gyrus (**b**) were imaged with confocal microscopy. Scale bars are 200 µm.

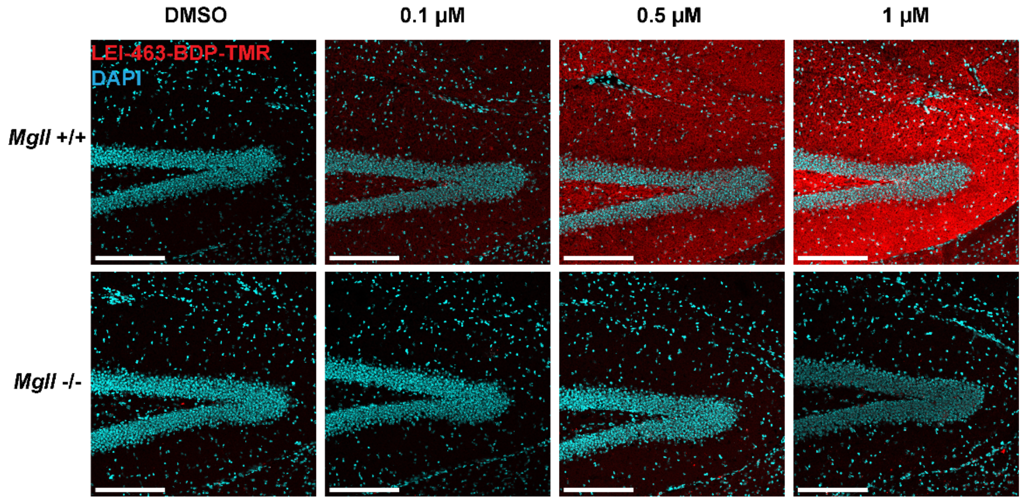


**Figure S4. Dose response visualization of MAGL with LEI-463-BDP-TMR. a**, Fresh-frozen cryosections from wildtype or *Mgll-/-* mouse brains were treated with LEI-463-BDP-TMR at the indicated concentrations (RT, 30 min), washed with THF/$H_2O$ (1/1) and subsequently the dentate gyrus was imaged with confocal microscopy. Scale bars are 200 µm.

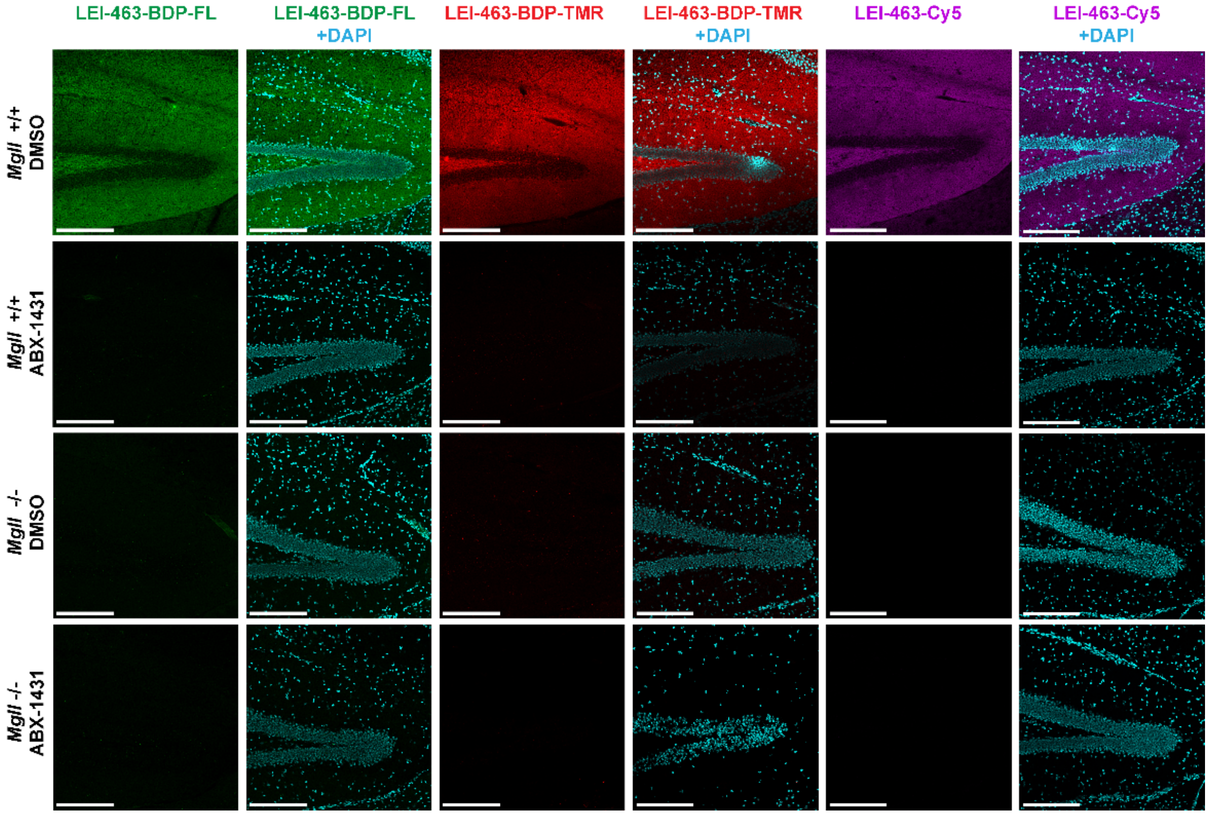


**Figure S5. LEI-463 probe signal is specific for MAGL. a**, Fresh-frozen cryosections from wildtype or *Mgll-/-* mouse brains were treated with vehicle or ABX-1431, and subsequently with the indicated LEI-463 probe (1 µM, RT, 30 min) and the hippocampal dentate gyrus was imaged with confocal microscopy. Scale bars are 200 µm.

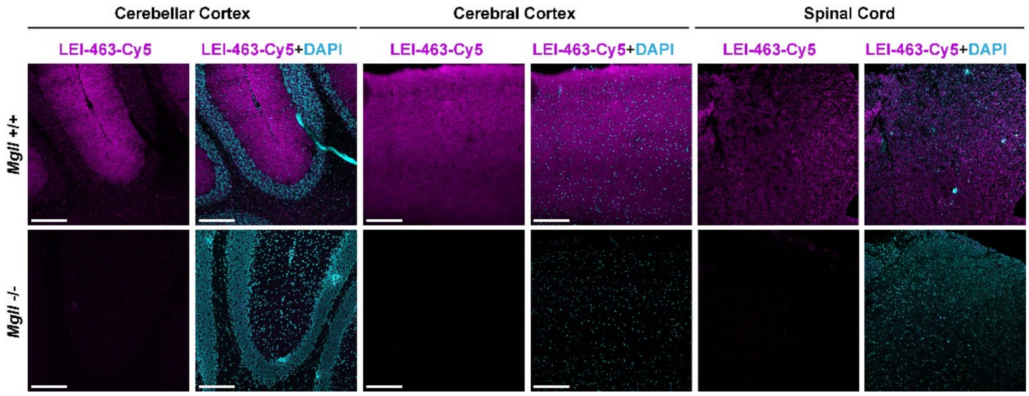


**Figure S6. Activity-based histology of MAGL in other CNS regions.** Fresh-frozen cryosections from wildtype or *Mgll*-/- mouse brains or spinal cords were treated with LEI-463 probe (1 μM, RT, 30 min) and subsequently imaged with confocal microscopy. Scale bars are 200 μm.

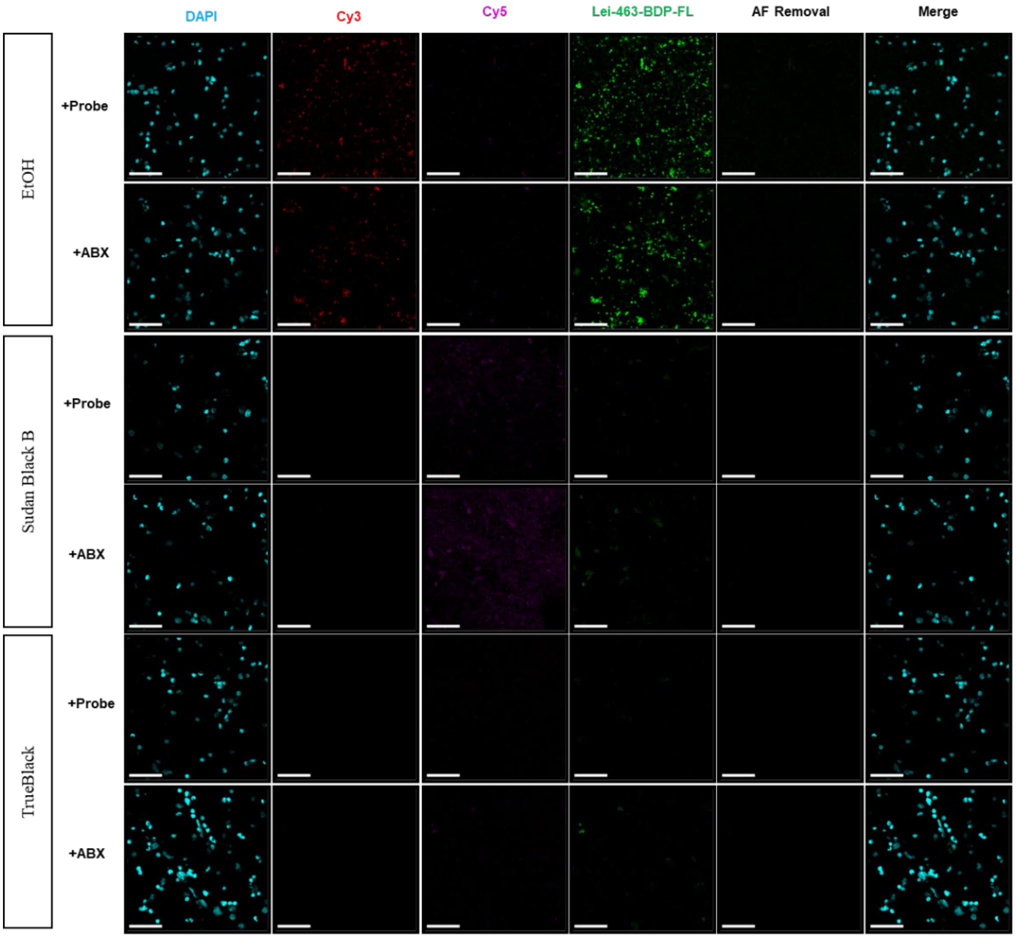


**Figure S7. Autofluorescence quenching using Sudan Black B and TrueBlack with MAGL probe LEI-463-BDP-FL.** Human cortical sections were treated with vehicle or ABX-1431, then treated with LEI-463-BDP-FL according to the general procedure, and lastly autofluorescence was quenched using Sudan Black B or Trueblack, compared to the 70% EtOH as vehicle. EtOH shows no AF quenching, while no probe signal was found. LEI-463-BDP-FL is outcompeted by AF signal and couldn't be visualized even after computational AF removal. Sudan Black quenches both Cy2 and Cy3 autofluorescence but not Cy5. Still no probe signal could be found. TrueBlack shows quenching of all AF channels, but again no probe signal could be retrieved. Scale bars are 50 µm.

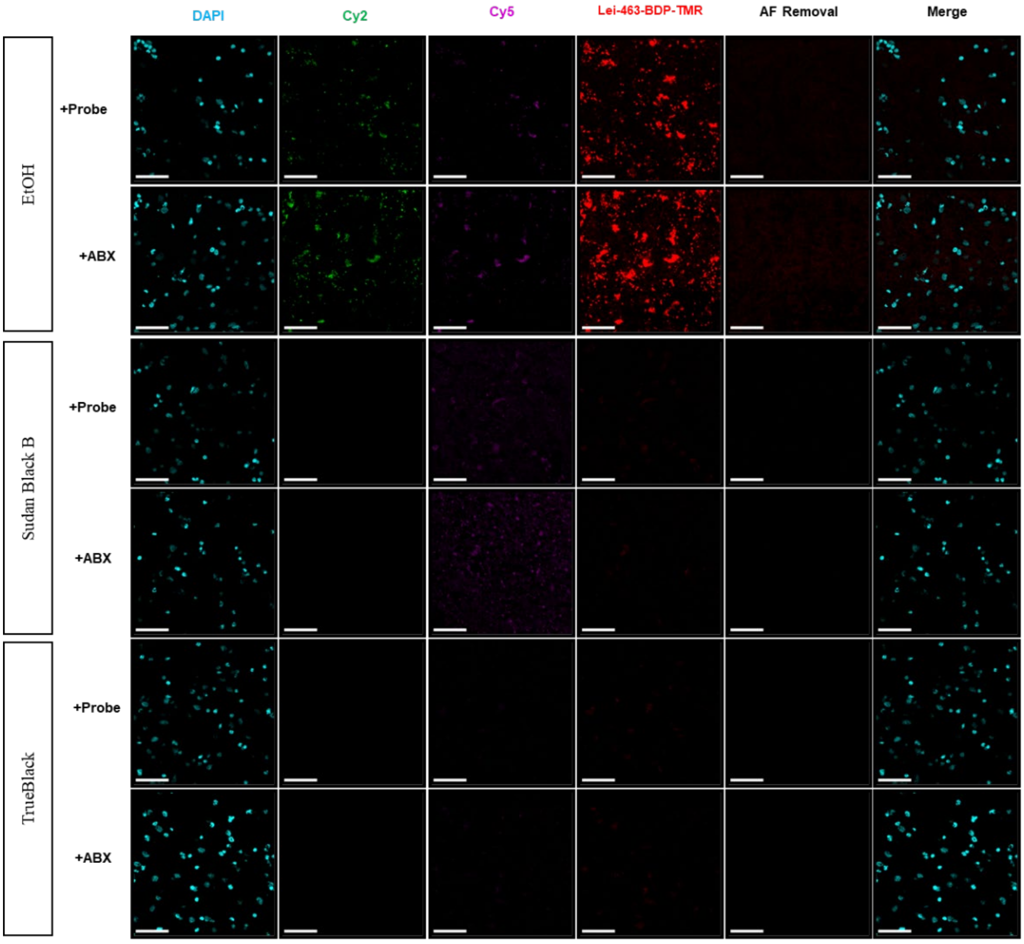


**Figure S8. Autofluorescence quenching using Sudan Black B and TrueBlack with MAGL probe LEI-463-BDP-TMR.** Human cortical sections were treated with vehicle or ABX-1431, then treated with LEI-463-BDP-TMR according to the general procedure, and lastly autofluorescence was quenched using Sudan Black B or Trueblack, compared to the 70% EtOH as vehicle. EtOH shows no AF quenching, while no probe signal was found. LEI-463-BDP-TMR is outcompeted by AF signal and couldn't be visualized even after computational AF removal. Sudan Black quenches both Cy2 and Cy3 autofluorescence but not Cy5. Still no probe signal could be found. TrueBlack shows quenching of all AF channels, but again no probe signal could be retrieved. Scale bars are 50 µm.

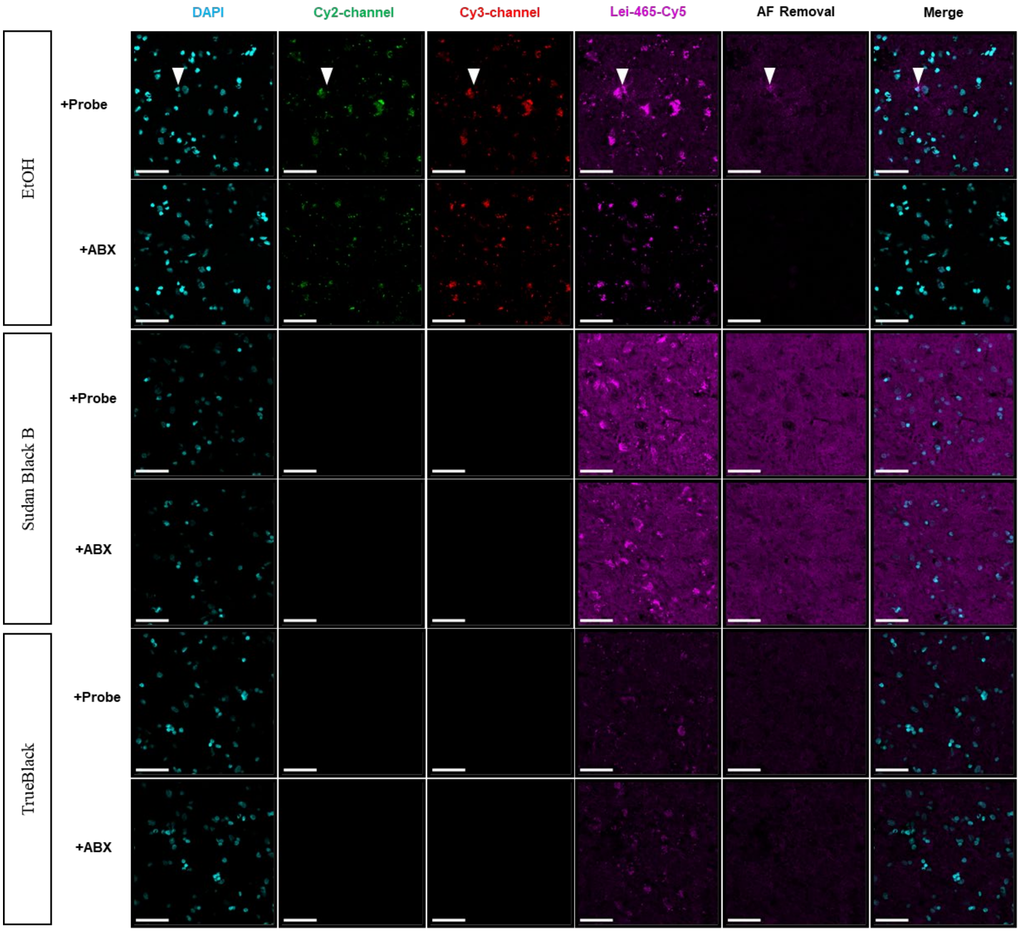


**Figure S9. Autofluorescence quenching using Sudan Black B and TrueBlack with MAGL probe LEI-463-Cy5.** Human cortical sections were treated with vehicle or ABX-1431, then treated with LEI-463-BDP-Cy5 according to the general procedure, and lastly autofluorescence was quenched using Sudan Black B or Trueblack, compared to the 70% EtOH as vehicle. Positive control using ethanol shows no AF quenching, but MAGL signal is found (arrowhead) which is blocked with ABX-1431. Sudan Black B and TrueBlack seem to introduce fluorescent background at increased laser powers, which ABX does not remove. Scale bars are 50 µm.

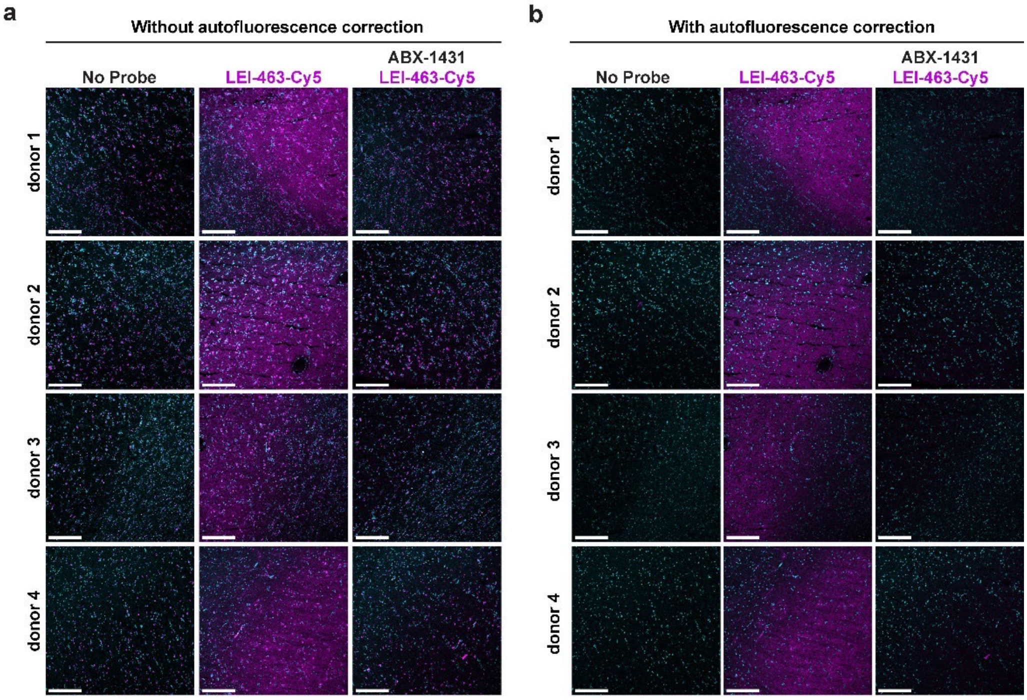


**Figure S10. LEI-463-Cy5 visualizes MAGL activity in human brains.** Fresh-frozen cryosections from cortical samples from four independent donors were treated with LEI-463 probe (1 µM, RT, 30 min) and subsequently imaged with confocal microscopy. **a**, Cy5 channel that was not corrected for autofluorescence. **b**, Cy5 channel that was corrected for autofluorescence using the Cy3 channel as an autofluorescence reference. Scale bars are 50 µm.

**Supplementary methods – organic synthesis**

**General remarks**

Reagents were purchased from Sigma Aldrich, Alfa Aesar, ChemScene, AnaSpec or ACROS organics at reagent grade and used without further purification. All moisture sensitive reactions were performed under a nitrogen atmosphere. Dry solvents were dried using 3 Å molecular sieves. Ethyl acetate was distilled before use. Glassware was oven dried (80 $^{0}$C) prior to use. $^{1}$H and $^{13}$C NMR spectra were recorded on a Bruker DPX-300 (300 MHz), AV-400 (400 MHz) or Bruker DRX-500 (500 MHz). Used software for interpretation of NMR-data was Bruker TopSpin 1.3 and MestReNova 11.0. Chemical shift values are reported in ppm in relation to tetramethylsilane as internal standard or solvent resonance as the internal standard ($CDCl_3$: δ 7.26 for $^{1}$H, δ 77.16 for $^{13}$C). Data are reported as follows: chemical shifts (δ), multiplicity (s = singlet, d = doublet, dd = double doublet, td = triple doublet, t = triplet, q = quartet, pentet = p, heptet = hept, br s = broad singlet, m = multiplet), coupling constants *J* (Hz), and integration (only for $^{1}$H-NMR). Liquid chromatography was performed on a Finnigan Surveyor LC/MS system, equipped with a C18 column. Flash chromatography was performed using SiliCycle silica gel type SiliaFlash P60 (230–400 mesh). TLC analysis was performed on Merck silica gel 60/Kieselguhr F254, 0.25 mm. Compounds were visualized using UV or $KMnO_4$ stain ($K_2CO_3$ (40 g), $KMnO_4$ (6 g), and water (600 mL)).

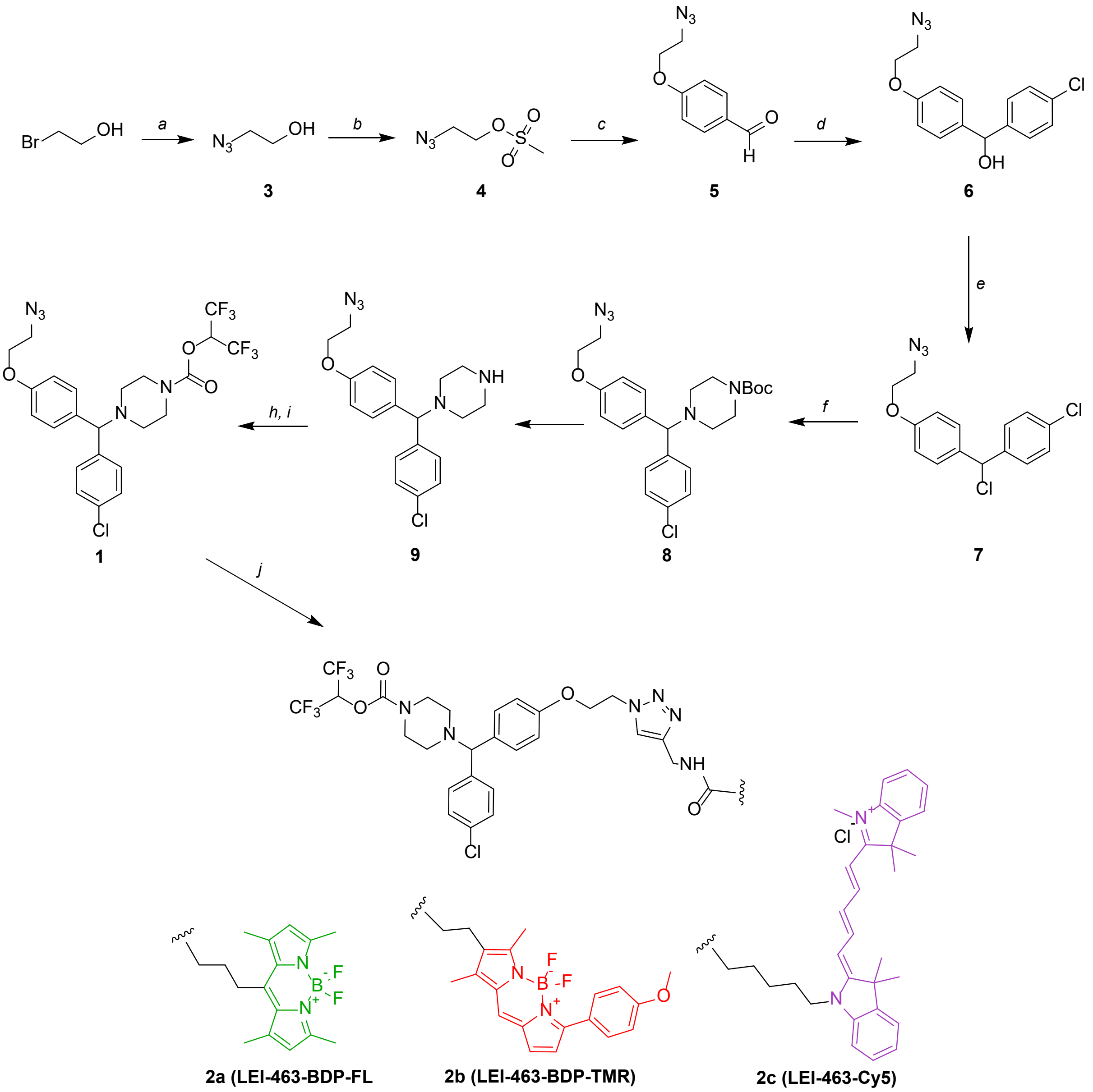

**Scheme 1. Synthesis of two-step ABPs targeting MAGL.** Reagents and conditions: *a*) sodium azide, $H_2O$, 80 °C, quantitative; *b*) mesyl chloride, DCM, 0-20 °C, quant.; *c*) 4-hydroxybenzaldehyde, $K_2CO_3$, DMF, 80 °C, 82%; *d*) 4-chlorophenyl magnesiumbromide, THF, -78 °C, 87%; *e*) thionyl chloride, DCM, 40 °C, quant.; *f*) *N*-Boc piperazine, $K_2CO_3$, DCM, 40 °C, 93%; *g*) HCl in dioxane, DCM, RT, 82%; *h*) triphosgene, $Na_2CO_3$, DCM, 0-20 °C; *i*, hexafluoroisopropanol, $Na_2CO_3$, DMF, RT, 47%. *j*, fluorophore-alkyne, $CuSO_4$, Sodium ascorbate, DCM/$H_2O$, RT, 50-93%.

**1,1,1,3,3,3-Hexafluoropropan-2-yl-4-((4-(2-azidoethoxy)phenyl)(4-chlorophenyl)methyl) piperazine-1-carboxylate (1)**

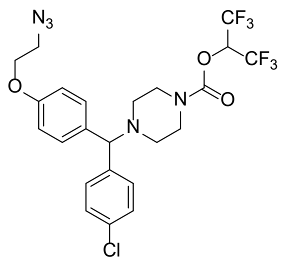

To a cooled (0 ˚C) solution of triphosgene (77 mg, 0.259 mmol) in dry DCM (5 mL) was slowly added under an inert atmosphere a solution of **7** (193 mg, 0.519 mmol) in dry DCM (2 mL). After the addition was completed, anhydrous $Na_2CO_3$ (110 mg, 1.038 mmol) was added and the solution was allowed to slowly warm to RT and stirred for 3 hours. The solution was filtered and the filtrate was evaporated to afford the carbamic chloride as a crude intermediate. This was dissolved in dry DMF (7 mL) and 1,1,1,3,3,3-hexafluoropropan-2-ol (0.273 ml, 2.59 mmol) and $Na_2CO_3$ (110 mg, 1.04 mmol) were added. The reaction was stirred under nitrogen atmosphere at RT for 3 hours and subsequently quenched with water and sat. aq. $NaHCO_3$. The mixture was diluted with water and the product extracted with DCM. The organic layer was dried ($MgSO_4$) and the solvent evaporated. The resulting crude product was purified over silica column chromatography (5-10% EtOAc in *n*-pentane) to afford **1** as a colourless thick oil (138 mg, 0.24 mmol, 47%). $^1$H NMR (400 MHz, $CDCl_3$) δ 7.33 (d, *J* = 8.5 Hz, 2H), 7.30 – 7.23 (m, 4H), 6.84 (d, *J* = 8.5 Hz, 2H), 5.74 (hept, *J* = 6.2 Hz, 1H), 4.20 (s, 1H), 4.10 (t, *J* = 5.1 Hz, 2H) 3.61 – 3.48 (m, 6H), 2.48 – 2.25 (m, 4H).$^{13}$C NMR (101 MHz, $CDCl_3$) δ 157.67, 151.47, 140.95, 134.20, 132.95, 129.05, 129.03, 128.98, 114.94, 74.48, 68.12 (hept*, *J* = 34.3 Hz), 67.10, 51.38, 51.22, 50.21, 44.94, 44.58. LC/MS: calculated for $[C_{23}H_{22}ClF_6N_5O_3 + H]^+$: 566.13, found: 565.60. * only 5 out of 7 peaks observed.

**1,1,1,3,3,3-Hexafluoropropan-2-yl 4-((4-chlorophenyl)(4-(2-(4-((4-(5,5-difluoro-1,3,7,9-tetramethyl-5*H*-4l4,5l4-dipyrrolo[1,2-c:2',1'-f][1,3,2]diazaborinin-10-yl)butanamido)methyl)-1*H*-1,2,3-triazol-1-yl)ethoxy)phenyl)methyl)piperazine-1-carboxylate (2a, LEI-463-BDP-FL)**

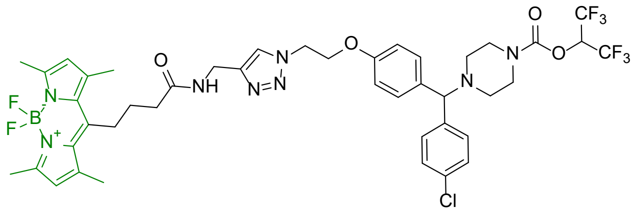

To a solution of **1** (35 mg, 62 µmol) in a 1:1 DCM/$H_2O$ suspension (2 mL) was added BODIPY 493/503 – alkyne (**13**, 22 mg, 59 µmol). Then, the mixture was purged with nitrogen and added were anhydrous $CuSO_4$ (9.5 mg, 31 µmol, 0.5 eq) and sodium-ascorbate (11.7 mg, 31 µmol, 0.5 eq). The reaction was vigorously stirred at RT overnight under nitrogen atmosphere. Subsequently, the reaction was diluted with $H_2O$ and extracted with DCM. The organic layer was dried ($MgSO_4$), filtrated and evaporated. The resulting crude was purified on silica column chromatography (0 – 10% MeOH in DCM) to afford **2a (LEI-463-BDP-FL)** as an orange solid (49 mg, 52 µmol, 88%). $^1$H NMR (400 MHz, $CDCl_3$) δ 7.70 (s, 1H), 7.29 (d, *J* = 8.5 Hz, 2H), 7.23 (d, *J* = 8.6 Hz, 4H), 6.76 (d, *J* = 8.7 Hz, 2H), 6.68 (t, *J* = 5.7 Hz, 1H), 6.02 (s, 2H), 5.72 (hept, *J* = 6.2 Hz, 1H), 4.68 (t, *J* = 4.8 Hz, 2H), 4.46 (d, *J* = 5.6 Hz, 2H), 4.27 (t, *J* = 5.0 Hz, 2H), 4.17 (s, 1H), 3.55 – 3.49 (m, 4H), 2.99 – 2.90 (m, 2H), 2.49 (s, 6H), 2.35 (m, =, 10H), 1.99 – 1.88 (m, 2H), 1.87 – 1.80 (m, 2H). $^{13}$C NMR (101 MHz, $CDCl_3$) δ 171.82, 157.10, 154.17, 151.46, 145.36, 140.82, 140.61, 134.85, 133.00, 129.09, 129.01, 129.00, 121.87, 114.30, 73.68, 68.09, 66.71, 51.34, 51.17, 49.33, 44.89, 44.54, 35.79, 33.99, 27.64, 27.42, 16.45, 14.57. HRMS: calculated for $[C_{43}H_{46}BClF_8N_8O_4 + H]^+$: 937.3376, found: 937.3372

**1,1,1,3,3,3-Hexafluoropropan-2-yl 4-((4-chlorophenyl)(4-(2-(4-((3-(5,5-difluoro-7-(4-methoxyphenyl)-1,3-dimethyl-5*H*-5l4,6l4-dipyrrolo[1,2-c:2',1'-f][1,3,2]diazaborinin-2-yl)propanamido)methyl)-1*H*-1,2,3-triazol-1-yl)ethoxy)phenyl)methyl)piperazine-1-carboxylate (2b, LEI-463-BDP-TMR)**

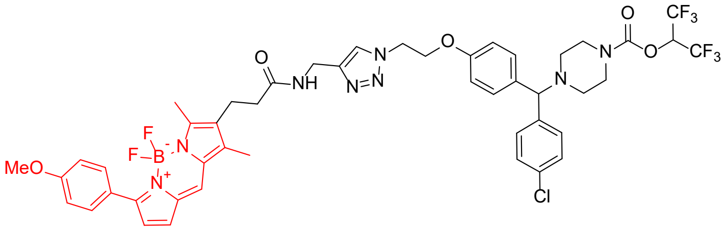

To a solution of **1** (8.7 mg, 15 µmol) in a 1:1 DCM/$H_2O$ suspension (2 mL) was added BODIPY-TMR – alkyne (Lumiprobe) (6.7 mg, 15 µmol). Then, the mixture was purged with nitrogen and added were anhydrous $CuSO_4$ (2.4 mg, 7.5 µmol, 0.5 eq) and sodium-ascorbate (3.0 mg, 7.5 µmol, 0.5 eq). The reaction was vigorously stirred at RT overnight under nitrogen atmosphere. Subsequently, the reaction was diluted with $H_2O$ and extracted with DCM. The organic layer was dried ($MgSO_4$), filtrated and evaporated. The resulting crude was purified on silica column chromatography (0 – 10% MeOH in DCM) to afford **2b (LEI-463-BDP-TMR)** as an dark red solid (11 mg, 11 µmol, 72%). $^1$H NMR (400 MHz, $CDCl_3$) δ 7.87 (d, *J* = 9.0 Hz, 2H), 7.57 (s, 1H), 7.30 (d, *J* = 8.4 Hz, 2H), 7.26 – 7.19 (m, 4H), 7.05 (s, 1H), 7.00 – 6.91 (m, 3H), 6.74 (t, *J* = 8.8 Hz, 2H), 6.53 (d, *J* = 4.1 Hz, 1H), 6.31 (t, *J* = 5.7 Hz, 1H), 5.72 (hept, *J* = 6.2 Hz, 1H), 4.57 (t, *J* = 5.1 Hz, 2H), 4.44 (d, *J* = 5.6 Hz, 2H), 4.22 – 4.14 (m, 3H), 3.84 (s, 3H), 3.57 – 3.46 (m, 4H), 2.73 (t, *J* = 7.5 Hz, 2H), 2.49 (s, 3H), 2.42 – 2.32 (m, 4H), 2.30 (d, *J* = 7.2 Hz, 2H), 2.16 (s, 3H). $^{13}$C NMR (101 MHz, $CDCl_3$) δ 170.61, 161.45, 156.52, 151.82, 145.17, 130.74, 128.99, 128.95, 128.03, 123.15, 122.86, 114.80, 113.81, 74.37, 73.23, 66.81, 55.32, 53.97, 51.32, 49.63, 45.36, 44.11, 36.20, 34.81, 19.45, 9.67. HRMS: calculated for $[C_{47}H_{46}BClF_8N_8O_5 + H]^+$: 1001.3326, found: 1001.3322

**2-((1*E*,3*E*)-5-((*E*)-1-(5-(((1-(2-(4-((4-Chlorophenyl)(4-(((1,1,1,3,3,3-hexafluoropropan-2-yl)oxy)carbonyl)piperazin-1-yl)methyl)phenoxy)ethyl)-1*H*-1,2,3-triazol-4-yl)methyl)amino)-5-oxopentyl)-3,3-dimethylindolin-2-ylidene)penta-1,3-dien-1-yl)-1,3,3-trimethyl-3H-indol-1-ium (2c, LEI-463-Cy5)**

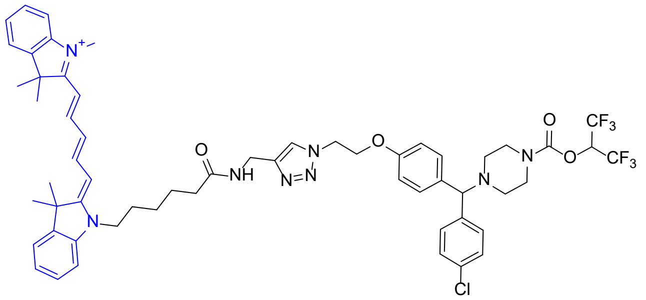

Compound **1** (25 mg, 44 µmol) was dissolved together with Cy5-alkyne (**14**, 23 mg, 44 µmol) in a nitrogen-purged 1:1 DCM/$H_2O$ suspension (2 mL). Then, anhydrous $CuSO_4$ (7.0 mg, 22 µmol, 0.5 eq) and sodium-ascorbate (8.7 mg, 22 µmol, 0.5 eq) were added and reaction was vigorously stirred at RT overnight under nitrogen atmosphere. Subsequently, the reaction was diluted with $H_2O$ and extracted with DCM. The organic layer was dried ($MgSO_4$), filtrated and evaporated. The resulting crude was purified on silica column chromatography (0 – 10% MeOH in DCM) to afford **2c (LEI-463-Cy5)** as a dark blue solid (24 mg, 22 µmol, 50%). A layer of NaCl (~5 g) was added on top of the silica to ensure elution of the chloride salt of **2c**. $^1$H NMR (400 MHz, $CDCl_3$) δ 8.43 (s, 1H), 8.03 (s, 1H), 7.86 (td, *J* = 13.0, 7.5 Hz, 2H), 7.41 – 7.28 (m, 6H), 7.26 – 7.19 (m, 6H), 7.11 (d, *J* = 8.0 Hz, 1H), 7.06 (d, *J* = 7.9 Hz, 1H), 6.80 (d, *J* = 8.5 Hz, 2H), 6.46 (d, *J* = 13.7 Hz, 1H), 6.28 (d, *J* = 13.5 Hz, 1H), 5.71 (hept, *J* = 6.2 Hz, 1H), 4.67 (t, *J* = 5.4 Hz, 2H), 4.55 (d, *J* = 5.1 Hz, 2H), 4.29 (t, *J* = 5.4 Hz, 2H), 4.17 (s, 1H), 4.06 (t, *J* = 7.6 Hz, 2H), 3.59 (s, 3H), 3.53 (m, 4H), 2.45 – 2.31 (m, 6H), 1.89 – 1.72 (m, 4H), 1.69 (s, 6H), 1.68 (s, 6H), 1.52 (m, 2H). HRMS: calculated for $[C_{58}H_{64}ClF_6N_8O_4]^+$: 1085.4634, found: 1085.4638 *amide proton not found*

**2-Azidoethanol (3)**

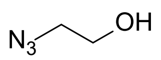


To a stirred solution of sodium azide (5.30 g, 82.5 mmol) in water (30 mL) was added 2-bromoethanol (2.3 mL, 33 mmol). The stirred was refluxed for 16 hours, and subsequently basified by addition of potassium hydroxide (1M, pH 13). The product was extracted with diethyl ether, the combined organic phases dried ($MgSO_4$) filtered and evaporated under reduced pressure to obtain **5** as a colourless liquid that was used without further purification (2.9 g, 33 mmol, quant.). $^{1}H$ NMR (400 MHz, $CDCl_3$) δ 3.82 – 3.68 (m, 3H), 3.41 (d, $J$ = 5.1 Hz, 2H). $^{13}C$ NMR (101 MHz, $CDCl_3$) δ 61.05, 53.29.

**2-Azidoethyl methane sulfonate (4)**

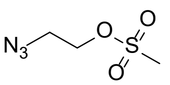


To a stirred and cooled (0 °C) solution of **3** (2.9 g, 33 mmol) in dichloromethane (30 mL) was dropwise added a solution of methanesulfonyl chloride in DCM (3.2 mL, 41.4 mmol) and subsequently slowly warmed up to RT. After stirring at RT for 2 hours, the reaction was quenched by the addition of water (15 mL), and the organic layer was dried ($MgSO_4$), filtered and evaporated to afford the crude product **4** as a yellow liquid, which was used without further purification (5.5 g, 33 mmol, quant). $^{1}H$ NMR (400 MHz, $CDCl_3$) δ 3.19 – 2.86 (m, 2H), 2.45 – 2.15 (m, 2H), 1.78 – 1.67 (m, 3H). $^{13}C$ NMR (101 MHz, $CDCl_3$) δ 67.85, 49.87, 37.69.

**4-(2-Azidoethoxy)benzaldehyde (5)**

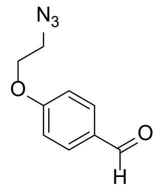


To a stirred solution of **4** (3.5 g, 21 mmol) in dimethylformamide (60 mL) were added 4-hydroxybenzalhyde (2.05 g, 16.8 mmol) and $K_2CO_3$ (3.5 g, 25 mmol). The reaction was vigorously stirred and heated up to 80 °C for 16 h. Afterwards, water (50 mL) was added and the product was extracted with diethyl ether (3 x 30 mL). The combined organic layers were washed with brine, dried ($MgSO_4$) evaporated to obtain the crude product as an orange liquid. The product was purified with silica column chromatography (gradient 5-20 % ethyl acetate in *n*-pentane) to obtain **5** as slightly yellow oil (3.21 g, 13.8 mmol, 82%). $^{1}H$ NMR (400 MHz, $CDCl_3$) δ 9.90 (s, 1H), 7.85 (d, $J$ = 8.8 Hz, 2H), 7.03 (d, $J$ = 8.8 Hz, 2H), 4.23 (d, $J$ = 4.6 Hz, 2H), 3.65 (d, $J$ = 4.6 Hz, 2H). $^{13}C$ NMR (101 MHz, $CDCl_3$) δ 190.82, 163.18, 132.07, 130.49, 114.89, 67.27, 50.02.

**(4-(2-Azidoethoxy)phenyl), 4-chlorophenyl methanol (6)**

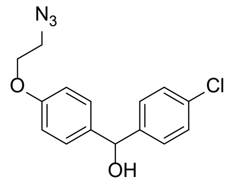


To a cooled (-78 °C) and stirred solution of **5** (1.2 g, 6.4 mmol) in dry tetrahydrofuran (60 mL) under nitrogen atmosphere was slowly added a solution of 4-chlorophenyl magnesium bromide (1M, 12.6 mL, 12.6 mmol) and subsequently stirred for 3 hours at -78 °C. After TLC analysis indicated full consumption of **5**, the reaction was quenched by the addition of saturated $NaHCO_3$ (10 mL). The mixture was extracted with DCM (3 x 50 mL) and the combined organic layers were washed with brine, dried ($MgSO_4$), filtered and evaporated to afford the crude product as a yellow oil. The product was purified over silica column chromatography (gradient 5-25 % ethyl acetate in *n*-pentane) to afford **6** as a thick colourless oil (1.73 g, 5.7 mmol, 87%). $^{1}H$ NMR (400 MHz, $CDCl_3$) δ 7.29 (s, 4H), 7.25 (d, $J$ = 8.8 Hz, 2H), 6.88 (d, $J$ = 8.7 Hz, 2H), 5.76 (d, $J$ = 2.8 Hz, 1H), 4.12 (t, $J$ = 4.7 Hz, 2H), 3.58 (t, $J$ = 4.9 Hz, 2H), 2.28 (d, $J$ = 2.9 Hz, 1H). $^{13}C$ NMR (101 MHz, $CDCl_3$) δ 157.97, 142.47, 136.60, 133.28, 128.68, 128.10, 127.87, 114.79, 75.21, 67.13, 50.83.

**_tert_-Butyl 4-((4-(2-azidoethoxy)phenyl)(4-chlorophenyl)methyl)piperazine-1-carboxylate (8)**

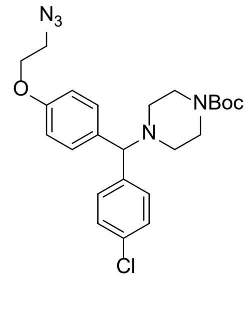


Under inert atmosphere thionyl chloride (1.6 ml, 21.92 mmol) was slowly added to solution of **6** (600 mg, 1.975 mmol) dry DCM (5 mL) and the reaction was heated up to 40 ˚C. After refluxing ON, the reaction was quenched with sat. aq. $NaHCO_3$ and the product was extracted using DCM. The combined organic layers were dried ($MgSO_4$), filtrated and evaporated to afford chlorinated compound **7** as a crude intermediate. Subsequently, the residue was dissolved in DCM (15 mL) and *tert*-butyl piperazine-1-carboxylate (1.1 g, 5.9 mmol) and potassium carbonate (1.4 g, 10 mmol) were added. The mixture was left to reflux overnight. The reaction was quenched through the addition of sat. aq. $NaHCO_3$ and water, and the product was extracted with DCM, dried ($MgSO_4$), filtered and evaporated. Purification was performed on silica column chromatography (5-20% EtOAc in *n*-pentane) to afford the title compound **8** as a slightly yellow thick oil (865 mg, 1.83 mmol, 93%) $^1$H NMR (400 MHz, $CDCl_3$) δ 7.34 (d, *J* = 8.2 Hz, 2H), 7.30 – 7.20 (m, 4H), 6.84 (d, *J* = 8.6 Hz, 2H), 4.17 (s, 1H), 4.09 (t, *J* = 4.4 Hz, 2H), 3.54 (t, *J* = 4.8 Hz, 2H), 3.41 (d, *J* = 4.9 Hz, 4H), 2.31 (d, *J* = 4.5 Hz, 4H), 1.43 (s, 9H). $^{13}$C NMR (101 MHz, $CDCl_3$) δ 157.46, 154.82, 142.27, 134.69, 132.63, 129.08, 129.02, 128.79, 114.75, 79.57, 74.61, 67.01, 60.43, 51.64, 50.15, 28.45. LC/MS: calculated for $[C_{24}H_{30}ClN_5O_3 + H]^+$: 471.20, found: 471.83

**1-((4-(2-Azidoethoxy)phenyl)(4-chlorophenyl)methyl)piperazine (9)**

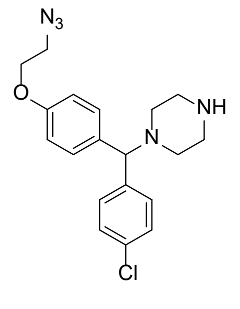


HCl in 1,4-dioxane (4M, 800 μL, 3.20 mmol) was added to solution of **8** (300 mg, 0.636 mmol) in EtOAc (2 mL). After stirring for 3 hours at RT, the reaction was quenched through the addition of sat. aq. $NaHCO_3$ and the product was extracted to EtOAc. The organic phase was then dried ($MgSO_4$) and evaporated. The product was purified by silica column chromatography (5-15% MeOH, 1% TEA in DCM) to afford title compound **9** as a slightly yellow thick oil (193 mg, 0.52 mmol, 82%). $^1$H NMR (400 MHz, $CDCl_3$) δ 7.31 (d, *J* = 8.5 Hz, 2H), 7.27 – 7.18 (m, 4H), 6.83 (d, *J* = 8.8 Hz, 2H), 4.09 (t, *J* = 4.9 Hz, 2H), 3.56 (t, *J* = 4.9 Hz, 2H), 3.26 – 3.15 (m, 4H), 2.75 – 2.54 (m, 4H). $^{13}$C NMR (101 MHz, $CDCl_3$) δ 157.18, 140.41, 133.72, 132.99, 129.01, 128.77, 128.75, 114.20, 74.01, 66.97, 50.07, 48.29, 45.97. LC/MS: $[C_{19}H_{22}ClN_5O + H]^+$ calculated: 371.15, found: 371.93

**Synthesis of clickable BODIPY-493/503 derivatives**

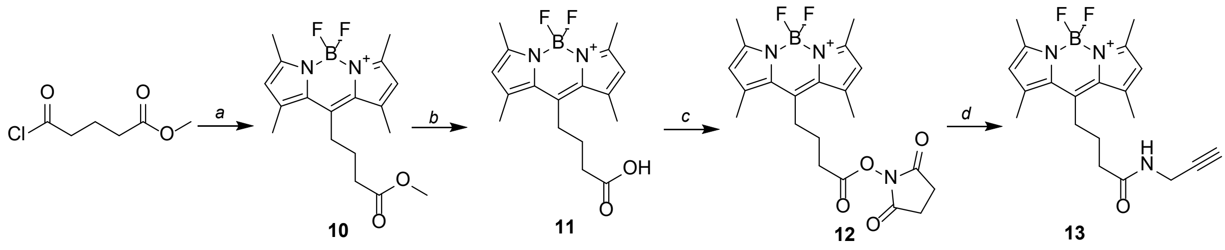


**Scheme 2. Synthesis of clickable BODIPY-493/503 derivatives.** *a)* 2,4-dimethyl-1*H*-pyrrole, TEA, $BF_3{\cdot}Et_2O$, DCM/toluene, 40 °C, 45%. *b)* NaOH, $H_2O$/THF, RT, quant. *c)* *N*-hydroxy succinimide, EDC, DCM, RT, quant. *d)* propargylamine, TEA, DCM, RT, 99%. *e)* 3-azidopropan-1-amine, TEA, DCM, 90%.

**Methyl-4-(5,5-difluoro-1,3,7,9-tetramethyl-5*H*-4l4,5l4-dipyrrolo[1,2-C:2',1'-f][1,3,2]diazaborinin-10-yl)butanoate (10)**

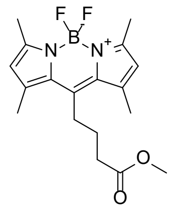

To a stirred solution of 2,4-dimethyl-1*H*-pyrrole (720 mg, 7.57 mmol) in DCM (30 mL) under nitrogen atmosphere was added methyl 5-chloro-5-oxopentanoate (0.7 mL, 5.07 mmol) and the mixture was refluxed for 2 hours. Successively, a solution of TEA (3.5 mL, 25.1 mmol) in toluene (20 mL) and $BF_3OEt_2$ (4.1 mL, 32.4 mmol) were added and the mixture was stirred at 50 $^{o}$C for another 2 hours. The reaction was quenched by the addition of saturated aqueous $NaHCO_3$. The product was extracted to DCM, washed with water, dried ($MgSO_4$), concentrated and subsequently purified by silica column chromatography (10-20% EtOAc in *n*-pentane), affording the title compound as a red/purple solid (600 mg, 1.7 mmol, 45%). $^{1}$H NMR (400 MHz, $CDCl_3$) δ 6.05 (s, 2H), 3.69 (s, 3H), 3.04 – 2.89 (m, 2H), 2.55 – 2.46 (m, 8H), 2.41 (s, 6H), 2.00 – 1.88 (m, 2H).

**4-(5,5-Difluoro-1,3,7,9-tetramethyl-5*H*-4l4,5l4-dipyrrolo[1,2-C:2',1'-f][1,3,2]diazaborinin-10-yl)butanoic acid (11)**

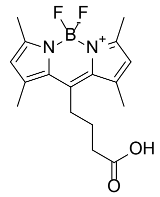

To a stirred solution of **10** (600 mg, 1,72 mmol) in THF (15 mL) was added aqueous sodium hydroxide (1.5 M, 12 mL, 18 mmol). After stirring overnight, the reaction was quenched by the addition of aqueous HCl until pH 3. The crude product was extracted with EtOAc and the solvents were removed under reduced pressure to afford the title compound as a purple solid, which was used without further purification (573 mg, 1.7 mmol, quant.). LC-MS (ESI+) m/z for $[C_{17}H_{21}BF_2N_2O_2 + H]^+$: calculated 334.17, found 335.08 (minor peak); m/z for $[C_{17}H_{21}BF_2N_2O_2 - F]^+$: calculated 315.17, found 315.25

**2,5-Dioxopyrrolidin-1-yl 4-(5,5-difluoro-1,3,7,9-tetramethyl-5*H*-4l4,5l4-dipyrrolo[1,2-C:2',1'-f][1,3,2]diazaborinin-10-yl)butanoate (12)**

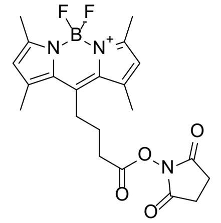

To a solution of **11** (573 mg, 1.72 mmol) in DCM (10 mL) were added *N*-hydroxy succinimide (296 mg, 2.57 mmol) and EDC (493 mg, 2.57 mmol). After 18 hours the reaction was quenched by the addition of water and the product extracted with DCM. The combined organic layers were dried ($MgSO_4$), filtrated and concentrated. Purification of the residue was performed on silica column chromatography (30%-70% EtOAc in *n*-pentane), affording **12** as a red solid (499 mg, 1.157 mmol, 100%). $^{1}$H NMR (400 MHz, $CDCl_3$) δ 6.06 (s, 2H), 3.17 – 3.05 (m, 2H), 2.93 – 2.83 (m, 4H), 2.81 (t, *J* = 7.0 Hz, 2H), 2.52 (s, 6H), 2.42 (s, 6H), 2.12 – 2.00 (m, 2H). LC-MS (ESI+) m/z for $[C_{21}H_{24}BF_2N_3O_4 + H]^+$: calculated 431.18, found 431.93 (minor peak); m/z for $[C_{21}H_{24}BF_2N_3O_4 - F]^+$: calculated 412.18, found 412.20

**4-(5,5-Difluoro-1,3,7,9-tetramethyl-5*H*-4l4,5l4-dipyrrolo[1,2-C:2',1'-f][1,3,2]diazaborinin-10-yl)-*N*-(prop-2-yn-1-yl)butanamide (13)**

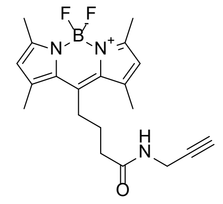

To a solution of **12** (104 mg, 0.241 mmol) in DCM were added propargylamine (20.1 µL, 0.314 mmol) and TEA (50.4 µl, 0.362 mmol). The reaction was stirred for 2 hours after which TLC analysis indicated full consumption of the starting material. The volatiles were evaporated and the residue purified by silica column chromatography (50% EtOAc in *n*-pentane) to afford the title compound as an orange solid (89 mg, 0.240 mmol, 99%) $^{1}$H NMR (400 MHz, $CDCl_3$) δ 6.05 (s, 2H), 5.65 (s, 1H), 4.13 – 3.95 (m, 2H), 3.14 – 2.89 (m, 2H), 2.51 (s, 6H), 2.44 – 2.27 (m, 9H), 2.24 (t, *J* = 2.6 Hz, 1H), 2.05 – 1.90 (m, 2H). LC-MS (ESI+) m/z for $[C_{20}H_{24}BF_2N_3O + H]^+$: calculated 371.20, found 317.13 (minor peak); m/z for $[C_{20}H_{24}BF_2N_3O - F]^+$: calculated 352.20, found 352.27 (majority)

**2-((1*E*,3*E*)-5-((*E*)-3,3-dimethyl-1-(6-oxo-6-(prop-2-yn-1-ylamino)hexyl)indolin-2-ylidene)penta-1,3-dien-1-yl)-1,3,3-trimethyl-3H-indol-1-ium (14)**

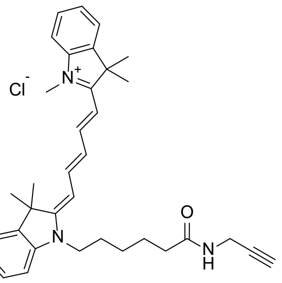


To a stirred solution of Cy5-carboxylic acid (80 mg, 0.15 mmol) in DCM (5 mL) was added perfluorophenyl trifluoroacetate (35 µL, 0.20 mmol) and DIPEA (70 µL, 0.4 mmol). After 3 hours, LC/MS analysis indicated full conversion of the starting material and excess of PFP-TFA was quenched by the adding $H_2O$ (18 µL). Subsequently, propargylamine (30 µL, 0.23 mmol) was added and the reaction was stirred for 18 hours. The mixture was diluted with water and extracted to DCM, the combined organic layers dried ($MgSO_4$), filtrated and evaporated. The compound was purified with silica gel column chromatography (1-5% MeOH in DCM). A layer of NaCl (~5 g) was added on top of the silica to ensure elution of the chloride salt of **26**. This afforded the title compound **26** (40 mg, 0.072 mmol, 47%). LC-MS (ESI+) m/z for $[C_{35}H_{42}N_3O]^+$: calculated 520.33, found 520.33. $^1H$ NMR (400 MHz, $CDCl_3$) δ 8.27 (s, 1H), 8.08 – 7.84 (m, 2H), 7.45 – 7.33 (m, 4H), 7.28 – 7.20 (m, 2H), 7.14 (d, *J* = 7.9 Hz, 1H), 7.10 (d, *J* = 7.9 Hz, 1H), 6.97 (t, *J* = 12.5 Hz, 1H), 6.57 (d, *J* = 13.6 Hz, 1H), 6.34 (d, *J* = 13.5 Hz, 1H), 4.11 (t, *J* = 7.7 Hz, 2H), 4.07 – 4.00 (m, 2H), 3.64 (s, 3H), 2.43 (t, *J* = 7.2 Hz, 2H), 2.13 (t, *J* = 2.5 Hz, 1H), 1.93 – 1.75 (m, 4H), 1.73 (s, 6H), 1.72 (s, 6H), 1.68 – 1.49 (m, 2H).
